# Severe COVID-19 shares a common neutrophil activation signature with other acute inflammatory states

**DOI:** 10.1101/2021.07.30.454529

**Authors:** Lena F. Schimke, Alexandre H.C. Marques, Gabriela Crispim Baiocchi, Caroline Aliane de Souza Prado, Dennyson Leandro M. Fonseca, Paula Paccielli Freire, Desirée Rodrigues Plaça, Igor Salerno Filgueiras, Ranieri Coelho Salgado, Gabriel Jansen-Marques, Antonio Edson Rocha Oliveira, Jean Pierre Schatzmann Peron, Gustavo Cabral-Miranda, José Alexandre Marzagão Barbuto, Niels Olsen Saraiva Camara, Vera Lúcia Garcia Calich, Hans D. Ochs, Antonio Condino-Neto, Katherine A. Overmyer, Joshua J. Coon, Joseph Balnis, Ariel Jaitovich, Jonas Schulte-Schrepping, Thomas Ulas, Joachim L. Schultze, Helder I. Nakaya, Igor Jurisica, Otávio Cabral-Marques

## Abstract

Severe COVID-19 patients present a clinical and laboratory overlap with other hyperinflammatory conditions such as hemophagocytic lymphohistiocytosis (HLH). However, the underlying mechanisms of these conditions remain to be explored. Here, we investigated the transcriptome of 1596 individuals, including patients with COVID-19 in comparison to healthy controls, other acute inflammatory states (HLH, multisystem inflammatory syndrome in children [MIS-C], Kawasaky disease [KD]), and different respiratory infections (seasonal coronavirus, influenza, bacterial pneumonia). We observed that COVID-19 and HLH share immunological pathways (cytokine/chemokine signaling and neutrophil-mediated immune responses), including gene signatures that stratify COVID-19 patients admitted to the intensive care unit (ICU) and COVID-19_nonICU patients. Of note, among the common differentially expressed genes (DEG), there is a cluster of neutrophil-associated genes that reflects a generalized hyperinflamatory state since it is also dysregulated in patients with KD and bacterial pneumonia. These genes are dysregulated at protein level across several COVID-19 studies and form an interconnected network with differentially expressed plasma proteins that point to neutrophil hyperactivation in COVID-19 patients admitted to the intensive care unit. scRNAseq analysis indicated that these genes are specifically upregulated across different leukocyte populations, including lymphocyte subsets and immature neutrophils. Artificial intelligence modeling confirmed the strong association of these genes with COVID-19 severity. Thus, our work indicates putative therapeutic pathways for intervention.

## 1. Introduction

During almost two years of Coronavirus disease 2019 (COVID-19) pandemic caused by the severe acute respiratory syndrome Coronavirus (SARS-CoV)-2, over 396 million cases and 5,7 million deaths have been reported worldwide (February 8^th^, 2022, WHO COVID-19 Dashboard). The clinical presentation ranges from asymptomatic to severe disease manifesting as pneumonia, acute respiratory distress syndrome (ARDS), and a life-threatening hyperinflammatory syndrome associated with excessive cytokine release (hypercytokinemia) [1–3]. Risk factors for severe manifestation and higher mortality include old age as well as hypertension, obesity, and diabetes [4]. Currently, COVID-19 continues to spread, new variants of SARS-CoV-2 have been reported and the number of infections resulting in death of young individuals with no comorbidities has increased the mortality rates among the young population [5,6]. In addition, some novel SARS-CoV-2 variants of concern appear to escape neutralization by vaccine-induced humoral immunity [7]. Thus, the need for a better understanding of the immunopathologic mechanisms associated with severe SARS-CoV-2 infection.

Patients with severe COVID-19 have systemic dysregulation of both innate and adaptive immune responses. In addition to highly activated CD4+ T cells[8] and high levels of autoantibodies linked to classic autoimmune diseases [9,10], they present with higher plasma levels of numerous cytokines and chemokines such as granulocyte macrophage colony-stimulating factor (GM-CSF), tumor necrosis factor (TNF), interleukin (IL)-6, soluble IL-6R, IL-8 (CXC chemokine ligand 8 (CXCL8), IL-18, monocyte chemoattractant protein-1 (MCP-1/CC chemokine ligand 2 [CCL2]) [11–13] than patients with moderate or mild COVID-19 disease [14], suggesting a more generalized hyperinflammatory condition. Notably, the hyperinflammation in COVID-19 shares similarities with cytokine storm syndromes such as those triggered by sepsis, autoinflammatory disorders, and metabolic conditions [15–17]. For instance, some COVID-19 patients may develop hyperinflammatory conditions such as the multisystem inflammatory syndrome in children (MIS-C), Kawasaki disease, and a severe hyperinflammatory state resembling a hematopathologic condition called hemophagocytic lymphohistiocytosis (HLH) [18]. All of them are life-threatening progressive systemic hyperinflammatory disorders characterized by multi-organ involvement. For instance, HLH patients may develop fever flares, hepatosplenomegaly and cytopenias due to hemophagocytic activity in the bone marrow [18–20] or within peripheral lymphoid organs such as pulmonary lymph nodes and spleen. HLH is marked by aberrant activation of B and T lymphocytes and monocytes/macrophages, coagulopathy, hypotension, and ARDS.

Therefore, we sought to characterize key signaling pathways and gene signatures associated with this more generalized hyperinflammatory state that is characteristic for some patients with severe COVID-19. The present study represents a follow-up of a recent report from our group [21], in which we performed an integrative analysis of transcriptional alterations in respiratory airways and peripheral blood leukocytes. This approach successfully developed by our and other groups [22–24] demonstrated multi-tissue systemic effects of SARS-CoV-2 infection, providing insightful mechanisms of SARS-CoV-2 pathology and cellular targets for therapy [23].

We first compared the molecular overlap between patients with COVID-19 and those with HLH, defined the transcriptomic and proteomic signatures stratifying COVID-19 patients admitted to the intensive care unit (COVID-19_ICU), and then investigated the behavior of the resulting molecular signature in other inflammatory syndromes and infectious diseases, enrolling a total of 1596 individuals, whose high throughput data was publicly available (**Table 1** and **Figure 1**).

**Table 1.**
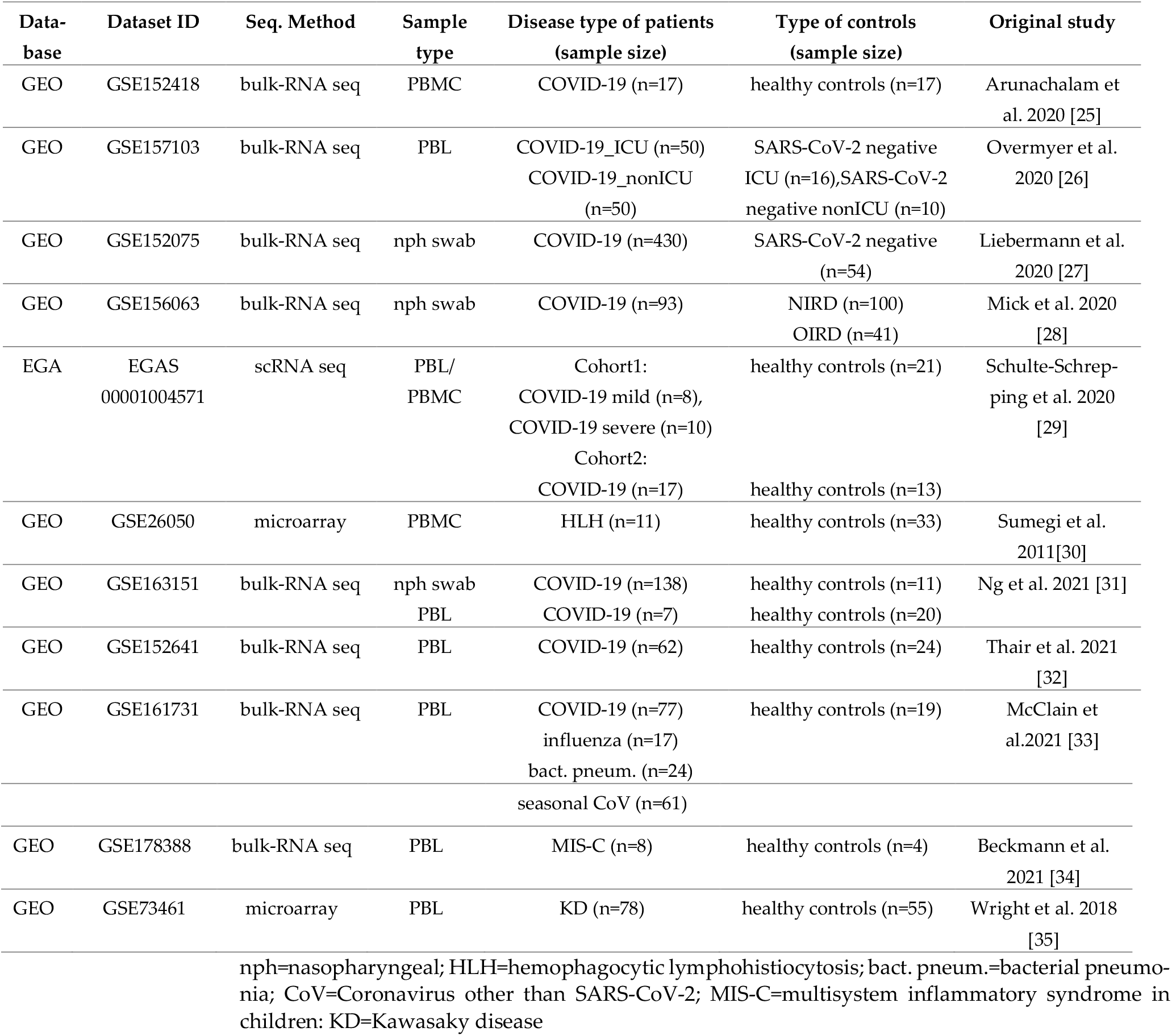
Dataset Information and sample size used for transcriptome analysis

**Figure 1.**
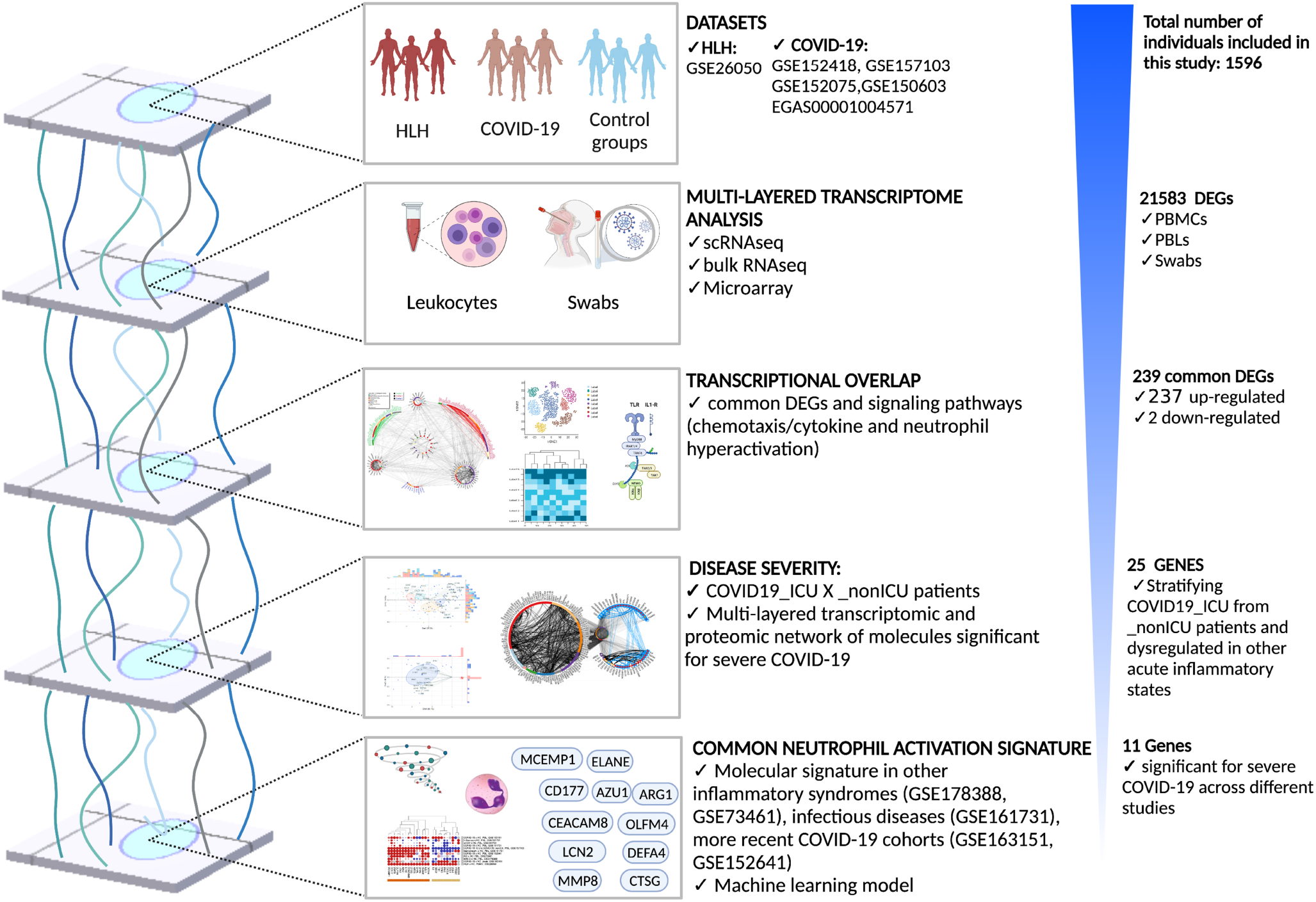
Schematic view of study workflow analyzed datasets and results obtained indicating common neutrophil activation signature between severe COVID-19 and other acute inflammatory states (Figure created using BioRender.com).

## 2. Materials and Methods

### Data Curation

We searched public functional genomics data repositories (Gene Expression Omnibus [36] and Array Express [37]) for human transcriptome data from patients with HLH and COVID-19 published until February 2021. During our analysis we included two more recently published COVID-19 datasets, two transcriptome studies from inflammatory diseases and one dataset with cohorts of other respiratory infectious diseases to compare with specific transcriptome signatures resulting from our first analysis. After evaluating study design, number of samples, and other relevant information (e.g., COVID-19 severity), we obtained raw count files (non-normalized) after trimming and alignment to the reference genome and followed guidelines to perform a meta-analysis report [38], which recomended to include at least three or four studies to reach a minimum of 1000 participants [39] in order to increase the statistical power of our analysis by increasing the signal-to-noise ratio. This resulted in a cohort of 1596 individuals derived from 11 datasets with transcriptome data generated from different platforms (**Table 1**).

### Differential expression analysis and visualization of transcriptional overlap

Read counts were transformed (log2 count per million or CPM) and differentially expressed genes (DEG) between groups were identified through the webtool NetworkAnalyst 3.0 [40] using limma-voom pipeline [41]. To determine DEGs of each dataset we applied the statistical cut-offs of log2 fold-change > 1 (up-regulated), log2 fold-change < −1 (down-regulated), and adjusted p-value < 0.05. Shared DEGs among all datasets were displayed using Venn diagram [42] and Circos Plot [43] online tools.

### Single cell RNAseq analysis

The Seurat Object containing the scRNAseq published by Schulte-Schrepping et al. [29] and deposited at the EGA (EGAS00001004571) were used for single cell analysis. We followed the Seurat pipeline [44] as previously described by Stuart et al. [45] to perform differential expression analysis and data visualization, i.e., UMAP, dotplot, and heatmap construction. Regression for the number of UMIs and scaling were performed as previously described [29].

### Interactome analysis

For a more comprehensive Protein–Protein Interaction (PPI) analysis, we used NAViGaTOR 3.0.14 [46] to visualize genes commonly dysregulated in COVID-19 and HLH datasets, highlighting the biological processes enriched by each gene. Prior to visualization, DEGs were used as input into Integrated Interactions Database (IID version 2021-05; http://ophid.utoronto.ca/iid) [47] to identify direct physical protein interactions. The resultant network was then annotated, analysed, and visualized using NAViGaTOR 3.0.14 [46]. The final figure was combined with legends using Adobe Illustrator ver. 26.0.3.

### Enrichment analysis and data visualization

We used ClusterProfiler [48] R package to obtain dot plots of enriched signaling pathways. Elsevier Pathway Collection analysis for selected gene lists (7 genes underlying HLH due to inborn errors of immunity (IEI) and 11 genes associated with severe COVID-19) was carried out using Enrichr webtool [49–51]. Sets of genes associated with cytokine/chemotaxis and neutrophil-mediated immunity from each dataset were visualized in bubble-based heat maps applying one minus cosine similarity using Morpheus [52]. Circular heatmaps were generated using R version 4.0.5 (The R Project for Statistical Computing. https://www.r-project.org/) and R studio Version 1.4.1106 (RStudio. https://www.rstudio.com/) using the circlize R package. Box plots were generated using the R packages ggpubr, lemon, and ggplot2.

### Correlation Analysis

Principal Component Analysis (PCA) of genes associated with COVID-19 severity (25 transcripts) was performed using the R functions prcomp and princomp, through factoextra package [53]. Canonical Correlation Analysis (CCA) [54] of genes associated with cytokines/chemokines and neutrophil-mediated immune responses was performed using the packages CCA and whitening [54]. In addition, we used the corrgram, psych, and inlmisc R packages to build correlograms. In addition, multilinear regression analysis for combinations of different variables (genes) was performed using the R package ggpubr, ggplot2 and ggExtra.

### Proteome Data Analysis

We also evaluated the proteomics data obtained from plasma samples of COVID-19 patients previously reported by Overmyer et al. [26]. Briefly, raw LFQ abundance values were quantified, normalized and log2 transformed as previously described [26]. Differences in protein expression between COVID-19_ICU and COVID-19_nonICU was calculated using the nonparametric MANOVA (multivariate analysis of variance) test [55] followed by analysis of nonparametric Inference for Multivariate Data [56] using the R packages npmv, nparcomp, and ggplot2. Enrichment of differentially expressed proteins (DEP) significant for COVID-19_ICU was performed using Enrichr webtool [49–51] and most significant enriched pathways were displayed by dot plot created with R using tidyverse, viridis and ggplot2 packages while Circos Plot of gene-pathway association was built using Circos online tool [43].

### Decision-tree classification and machine learning model predictors

We employed random forest model to construct a classifier able to discriminate between COVID-19_nonICU and COVID-19_ICU highlighting the most significant predictors for ICU admission. We trained a Random Forest model using the functionalities of the R package randomForest (version 4.6.14) [57]. Five thousand trees were used, and the number of variables resampled were equal to three. Follow-up analysis used the Gini decrease, number of nodes, and mean minimum depth as criteria to determine variable importance. Interaction between pairs of variables was assessed by minimum depth as criterion. The adequacy of the Random Forest model as a classifier was assessed through out of bags error rate and ROC curve. For cross-validation, we split the dataset in training and testing samples, using 75% of the observations for training and 25% for testing.

## 3. Results

### The transcriptional overlap between COVID-19 and HLH

We first performed a cross-tissue analysis of transcriptomic datasets obtained from peripheral blood lymphocytes (PBLs), peripheral blood mononuclear cells (PBMCs), and nasopharyngeal swabs. An association between the transcriptome information across the blood and respiratory airways of COVID-19 patients has been reported by our and other groups [21,23,24]. In this first approach, we obtained a total of 21,583 DEGs from seven COVID-19 cohorts from 5 datasets (both datasets GSE156063 and EGAS00001004571 have two different cohorts) and one HLH cohort (**Figure 2A** and **Supplementary Table S1**). Three other COVID-19 studies (GSE163151, GSE152641, and GSE 161731) were only included during our analysis because they were publicly available only after the beginning of our study. To identify the common DEGs we divided the datasets into three subgroups based on type of samples and RNAseq platforms: Overlap 1 (HLH and COVID-19 blood transcriptomes), Overlap 2 (HLH and COVID-19 nasopharyngeal swab transcriptomes), and Overlap 3 (HLH and COVID-19 scRNAseq transcriptomes) (**Supplementary Figure S1A and S1B**, and **Supplementary Table S2 and S3**). Even though the total number of DEGs of each dataset has large variability, the number of shared DEGs between the HLH and each COVID-19 dataset was similar across all studies and resulted in a total of 239 unique common DEGs between HLH and all COVID-19 datasets, most of them (237 DEGs) being up-regulated (**Figure 2B**). Hereafter, we focused on the implications of the up-regulated genes, since the 2 common down-regulated genes (granulysin or *GNLY*; myomesin 2 *or MYOM2*) alone did not enrich any significant pathway. However, this might also indicate a defect in cytotoxic activity, typical of HLH [58], that will require future investigation. The 237 common up-regulated DEGs encode proteins mainly involved in immune system, metabolic and signaling processes, forming a highly connected biological network based on physical protein-protein interactions (PPI, **Figure 2C**). Among them are important genes encoding molecules involved in activation of inflammatory immune responses (e.g., PGLYRP1, OLR1, FFAR2), cytokine and chemokine mediated immune pathways (e.g., IL1R2, CXCR2, CXCR8, CCL4, CCL2), and neutrophil activation (e.g., CD177, MPO, ELANE). Of note, the transcriptional overlap between HLH and COVID-19 contains several molecules interacting with 7 HLH/IEI-associated genes, which themselves were not among our DEGs (**Figure 2C**).

**Figure 2:**
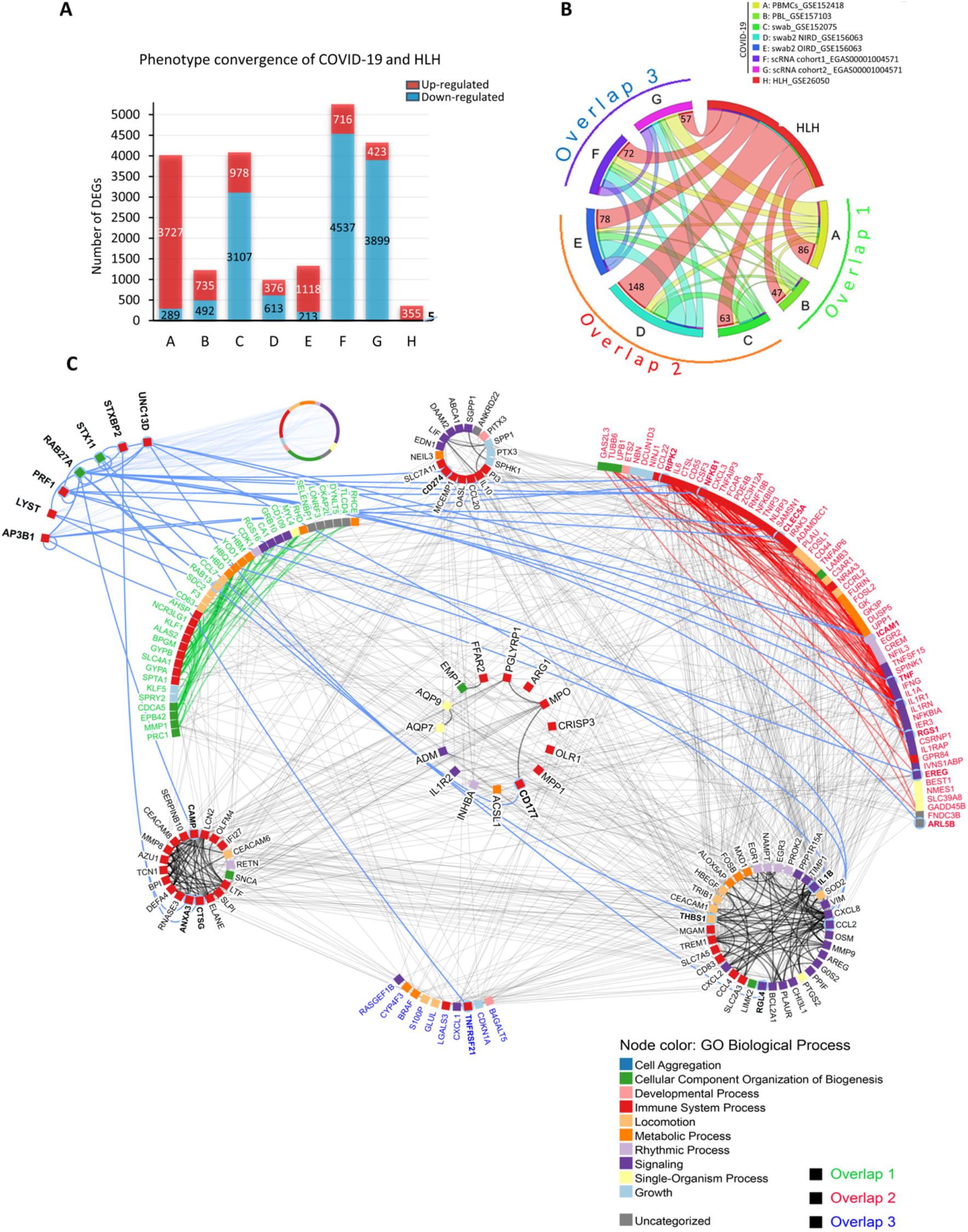
Transcriptional overlap between COVID-19 and HLH. (**A**) Number of differentially expressed genes (DEGs, up- and down-regulated) by dataset. (**B**) Circos plot showing 237 common up-regulated DEGs between HLH and the different COVID-19 datasets (red lines: number at the end of each line indicates exact number of shared DEGs), divided in 3 overlapping subgroups (Detailed in Supp. Table 2 and 3). The thickness of each line represents the number of genes shared between the different datasets. (**C**) Protein-protein interaction network among the 237 transcripts and 7 genes causing HLH due to inborn errors of immunity (IEI). Node colours denote Gene Ontology Biological Processes. The label (gene name) colours represent transcripts from *Overlap 1* (green), *Overlap 2* (red), and *Overlap 3* (blue). Center circle and side circles represent common molecules across all 3 or 2 overlapping datasets, respectively. The upper left subnetwork represents the interactions between the 7 genes associated with HLH and those from overlaps are bold. The circle on upper left (gene names not shown) contains 1329 proteins connected by 217 interactions with the 7 HLH/IEI-associated genes. The full network comprises 1538 proteins and 2522 direct physical interactions obtained from IID database ver. 2021-05 [47].

### Cytokine/chemotaxis and neutrophil signatures predominate in COVID-19 and HLH

We next dissected the biological functions enriched by the 237 common up-regulated DEGs between COVID-19 and HLH patients by performing enrichment analysis of biological processes (BPs) and cellular components (CCs) by these 237 DEGs. The top 20 most enriched BPs are demonstrated in **Figure 3A**, which encompass cytokine/chemotaxis and neutrophil-mediated innate immune responses, ranging from neutrophil activation, degranulation, and migration to responses to IL-1 as well as anti-microbial humoral response, (for all BPs see **Supplementary Table S4**). The CCs enriched (**Figure 3B**) include several compartments such as secretory granule lumen and membrane, azurophil tertiary and specific granules, as well as collagen-containing extracellular matrix, phagocytic vesicles, and primary lysosomes (**Supplementary Table S5**).

**Figure 3:**
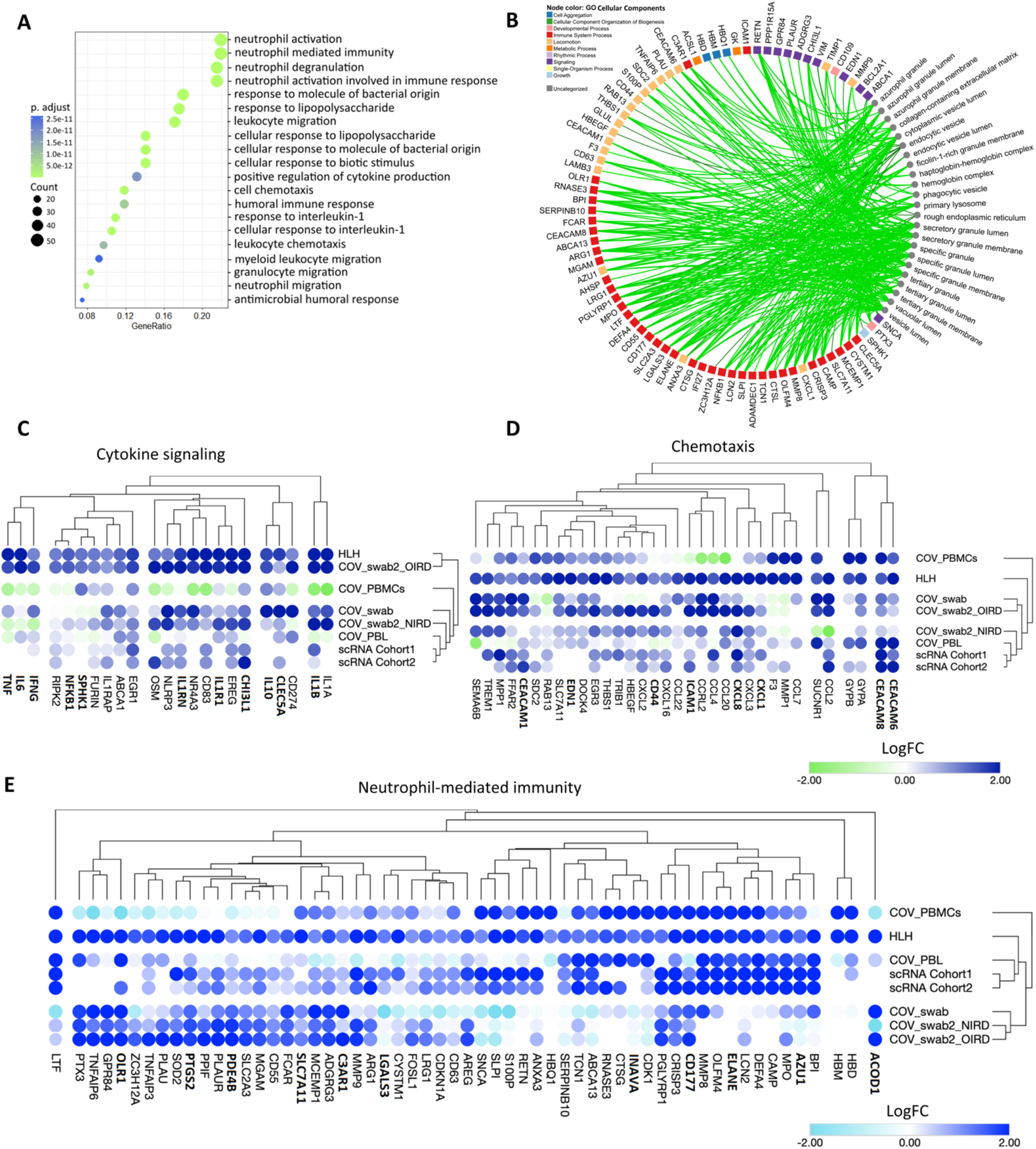
Cytokine/chemotaxis and neutrophil-associated transcriptional signatures predominate in the COVID-19 and HLH overlap. **(A)** Dot plot showing the most significant biological processes enriched by the 237 common up-regulated transcripts of COVID-19 and HLH datasets. The dot size is proportional to the number of genes enriching the ontology term and color proportional to adjusted p value (green > significant than blue). **(B)** Network highlighting genes and cellular component associations. Only enriched terms with adjusted *p* value <0.05 are shown by small grey circles. The degree of associations is displayed by edge color and thickness (e.g., lighter color and thinner edges signify fewer connections). Node color represents different GO CCs. Both enriched CCs and BPs were analyzed using ClusterProfiler with R programming. **(C-E)** Bubble heatmaps showing the hierarchical clustering based on Euclidian distance of expression patterns of genes associated to **(c)** cytokine signaling, **(D)** chemotaxis, and **(E)** neutrophil-mediated immunity in COVID-19 and HLH datasets. The color of circles corresponds to log2 fold change (log2FC). Pleiotropic genes belonging to more than one category are bold (**Supplementary Table S8**).

Of note, cytokine/chemotaxis and neutrophil signatures predominate in the COVID-19 and HLH multi-layered transcriptional overlap. A total of 25, 34, and 58 DEGs are assigned to cytokine, chemotaxis, and neutrophil signatures, respectively (**Figure 3C-E**: complete categorization can be seen in **Supplementary Table S6 and S7**). Several genes play pleiotropic roles in these gene ontology (GO) categories such as *CEACAM8, IL-1β, IL-6, EDN1, NFKB1* and *PDE4B* (**Supplementary Table S8**). For clarity in data visualization, we assigned these genes to a unique category (based on their predominant immunological function according to literature and GeneCards [59] or the human gene database). Among these are genes that code for chemokines and chemokine receptors that attract both lymphocytes and myelocytes to inflammation sites (CCL20, CCL2, CXCR1, CXCR3, CXCL8) [60,61], pro-inflammatory cytokines and cytokine receptors (IL-1B, IL-1R1, NFKB1, IFNG, IL-6, TNF) [62,63] that promote the activation of immune cells, and several proteins/granules with antimicrobial activity (MPO [64], AZU1 [65], ELANE [66,67], DEFA4 [68]). Moreover, there are metalloproteinases (MMP8 and MMP9) involved in degradation of extracellular matrix (ECM) to facilitate neutrophil migration [69,70] into the airways and in the regulation of cytokine activity. Of note, hierarchical clustering analysis of these genes indicated a cross-study grouping of closely functional-related molecules. For instance, *IFNG, IL6*, and *TNF; IL1A* and *IL1B*; as well as signaling molecules involved in the nuclear factor-κB (NF-κB) signaling such as *NFKB1, SPHK1*, and *RIPK2* clustered together in the cytokine group. Likewise, *GYPA* and *GYPB, CEACAM6* and *CEACAM8*, as well as *CCL* and *CXCL* chemokines in the chemotaxis group, and antimicrobial-related peptides such as *AZU1, MPO, CAMP, DEFA4, LCN2, ELANE, OLFM4*, and *CD177* in neutrophil-mediated immunity clustered together. However, we cannot exclude that this clustering pattern just represents a random tendency due to upstream GO categorization.

Relevant literature has emphasized that the 7 genes (*AP31B, LYST, PRF1, RAB27A, STX11, STXBP2, UNC13D*) known to cause fHLH (classically defined as familial HLH syndromes and hypopigmentation syndromes) [17,71] contribute to the dominant role played by T and NK cells in the development of HLH [72–74]. However, although these 7 genes are not commonly dysregulated across the datasets of COVID-19 and HLH patients, they also enrich several CCs (secretory vesicles, azurophilic granules or specific granules) and BPs (neutrophil degranulation) involved in the neutrophil immune response (**Supplementary Figure S2**). This result is in agreement with the role of these genes in a variety of neutrophil functions such as degranulation and formation of neutrophil extracellular traps (NET) [75–79].

### The relationship between cytokine/chemotaxis and neutrophil-mediated immunity gene signatures

Considering the potential association between cytokine/chemotaxis and neutrophil-mediated immunity representing regulatory and effector functions involved in the COVID-19 pathogenesis, we next analysed the relationship pattern and degree between these transcriptional signatures. We chose the COVID-19_PBL dataset provided by Overmyer et al. [26], which contains transcripts from 100 individuals with COVID-19 and 26 individuals with respiratory symptoms but negative for SARS-CoV-2, serving as control group (further explored in the next session). We performed canonical-correlation analysis (CCA), which is a multivariate statistical method to determine the linear relationship between two groups of variables [80]. In accordance with the cross-study hierarchical clustering, CCA revealed a strong association between several cytokine/chemotaxis related genes (e.g., *CXCL8, CEACAMs [1/6/8], IL1RAP, IL1R1, IL1B, NFKB1*) with those involved in neutrophil-mediated immune responses (e.g., *CTSG, ELANE, MMP8, TCN1*) in both COVID-19 patients and controls (**Figure 4A** and **4B**). Bivariate correlation analysis showed a similar phenomenon (**Supplementary Figure S3**). However, these correlation patterns partially changed when comparing COVID-19 with the control group. For instance, while reducing the correlation between molecules including *IL-10, CXCL8, NFKB1, ARG1*, and *SOD2*, new strong associations appeared between *ELANE, DEFA4, AZU1, CTSG*, and *LCN2*, with an overall tendency to higher relationships amid neutrophil-mediated immunity related genes in COVID-19 patients. **Figure 4C** illustrates this observation by scatter plots for some of these variables.

**Figure 4:**
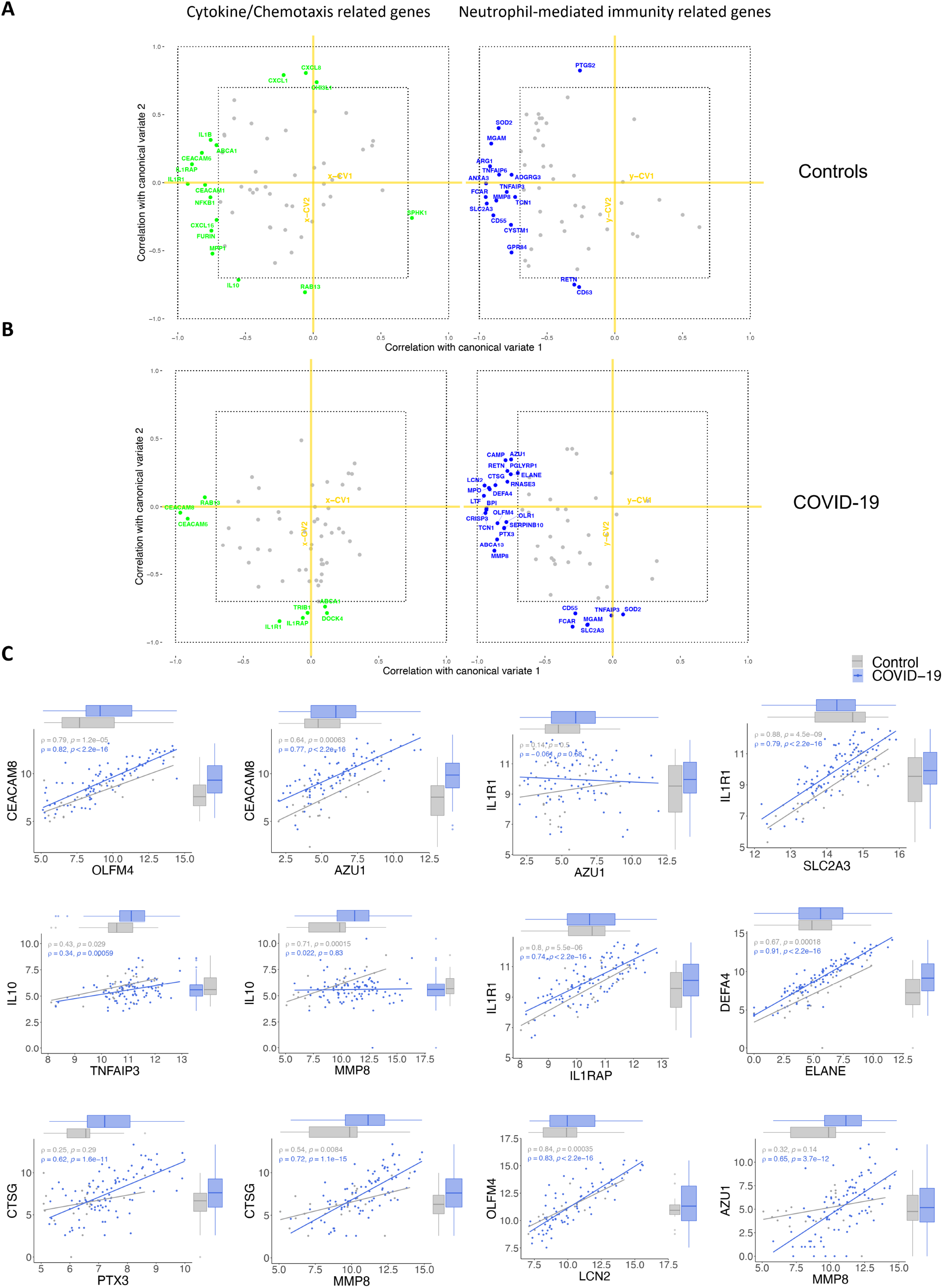
Infection with SARS-CoV-2 impacts the correlation between cytokine/chemotaxis and neutrophil-mediated immunity genes. (**A** and **B**) Estimated correlations of cytokine signaling/chemotaxis and neutrophil-mediated immunity molecules ranging from −1 to 1 versus their corresponding first 2 canonical variates (x-CV1 and x-CV2 for cytokine/chemotaxis related genes; y-CV1 and y-CV2 for neutrophil-mediated immunity genes) in (A) controls and (B) COVID-19 patients. Cytokine/chemotaxis and neutrophil-mediated immunity genes with a Spearman rank correlation of ≥ 0.7 are colored in green and blue, respectively, while those with a Spearman rank correlation of < 0.7 are gray in both groups. (**C**) Scatter plots with marginal boxplots display the relationship between variables (genes). Correlation coefficient (ρ) and significance level (p-value) for each correlation are shown within each graph.

### Transcripts stratifying severe COVID-19 from other respiratory diseases are highly dysregulated in HLH and other acute inflammatory states

Next, we assessed which genes of cytokine/chemotaxis signaling and neutrophil-mediated immune responses discriminate COVID-19 patients according to disease severity. We further investigated the COVID-19_PBL dataset (GSE157103) [26] comparing COVID-19 patients admitted to the intensive care unit (COVID-19_ICU) with those admitted to non-ICU units (COVID-19_nonICU). The severity of COVID-19 patients at ICU admission was defined based on APACHE II and SOFA scores [81] according to Overmyer et al. [26] (**Figure 5A**). Among all genes, 25 (15 up-regulated and 10 down-regulated genes) were differentially expressed between COVID-19_ICU and COVID-19_nonICU patients (**Supplementary Table S9**). Of note, most of these 25 genes have also been identified at protein level as dysregulated in COVID-19 patients across different studies (published during the development of our study; **Supplementary Table S10**). In addition, these 25 genes seem to belong to a systemic immune network of molecules induced by SARS-CoV-2 since they are also highly interconnected with 158 proteins (Supp. Table 11) significantly dysregulated in the plasma of COVID-19_ICU when compared to COVID-19_nonICU patients. Thus, they show several interactions and functional overlap (**Figure 5B**) with plasma proteins involved in neutrophil degranulation and neutrophil-mediated immunity (**Supplementary Figure S4** and **Supplementary Table S12**).

**Figure 5:**
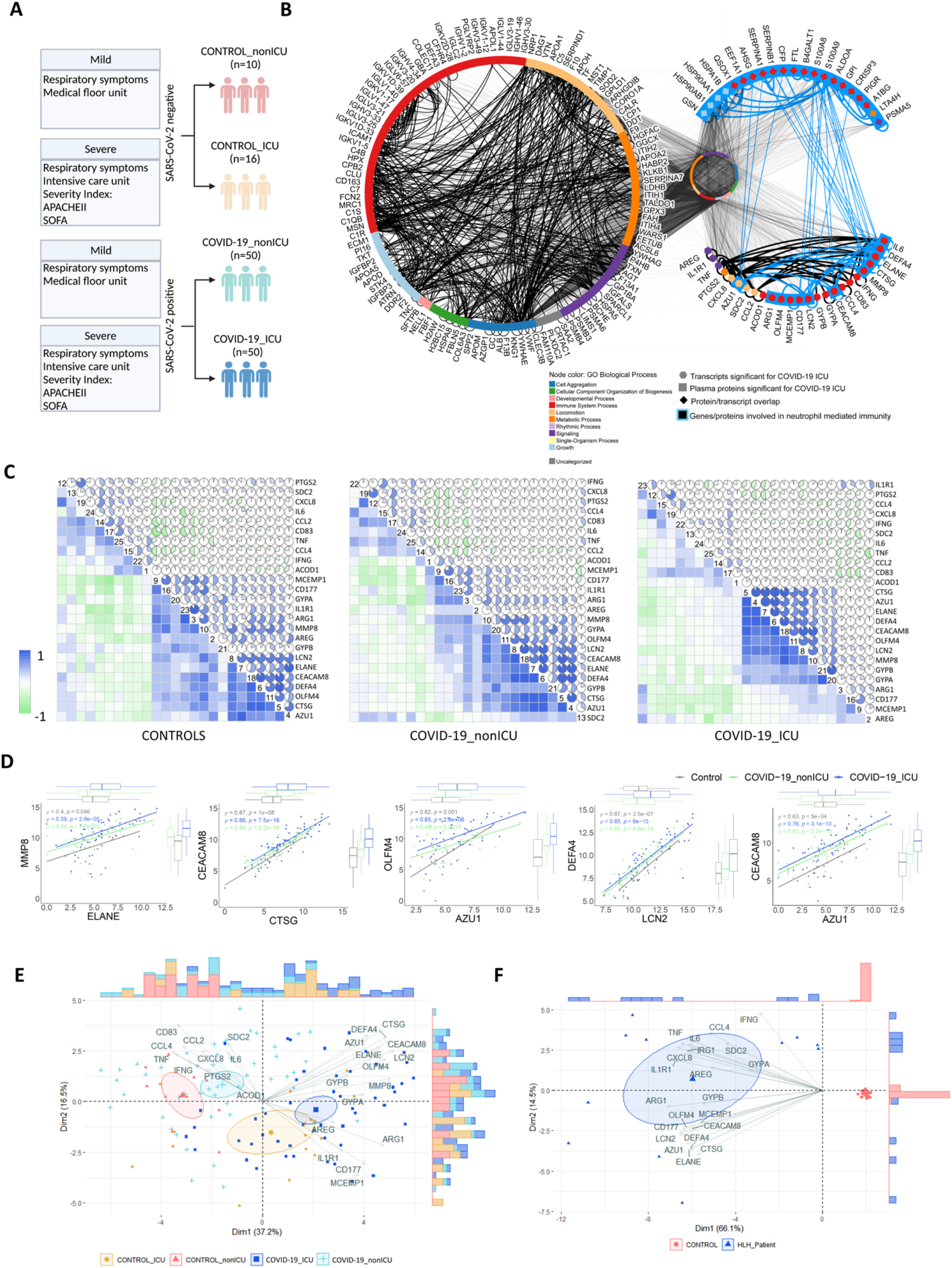
Transcripts stratifying severe COVID-19 from other respiratory diseases and HLH from healthy controls. **(A)** Schematic overview of study design and patient classification of dataset GSE157103 reported by Overmyer et al. [26]. **(B)** Protein-protein interaction (PPI) network highlighting interactions among the 158 proteins and the 25 genes significant for severe COVID-19_ICU) while keeping their other interacting partners (n=9,921) in the middle circle. Node colour denotes Gene Ontology Biological Process terms. Left circle shows 123 proteins and 554 interactions, upper right half circle shows 21 proteins and 29 interactions, and lower right side half circle shows 25 proteins and 65 interactions. Molecules involved in neutrophil-mediated immunity are highlighted with blue node outline. **(C)** Correlation matrices of the 25 DEGs (Controls, left matrix; COVID-19_nonICU, middle matrix; and COVID-19_ICU, right matrix). The color scale bar represents the Pearson’s correlation coefficient, containing negative and positive correlations from −1 to 1, respectively. **(D)** Scatter plots with marginal boxplots display the relationship between the eight genes stratifying severe COVID-19. Correlation coefficient (ρ) and significance level (p-value) for each correlation are shown within each graph. **(E)** Principal Component Analysis (PCA) with spectral decomposition shows the stratification of COVID-19_ICU from COVID-19_nonICU and other respiratory diseases (Control_nonICU and Control_ICU). Variables with positive correlation are pointing to the same side of the plot, contrasting with negative correlated variables, which point to opposite sides. Confidence ellipses are shown for each group/category. Bar plots associated with the PCA represent the sample distribution across the PCA axes. **(F)** PCA displaying the stratification of HLH and healthy controls based on the same 25 DEGs as in (E).

Bivariate correlation analysis based on these 25 genes showed that while controls and COVID-19_nonICU patients have a similar general cluster distribution, COVID-19_ICU patients tend to differ revealing only 8 genes with high positive correlations (**Figure 5C** and **Figure 5D**). To investigate the stratification power of these 25 DEGs, we performed principal component analysis (PCA) using a spectral decomposition approach [82,83], which examines the covariances/correlations between variables. This approach revealed that these DEGs clearly divide COVID-19_ICU, COVID-19_nonICU, Control_ICU and Control_nonICU (due to other respiratory illness but negative for SARS-CoV-2) groups (**Figure 5E** and **Supplementary Figure S5A** and **S5C**). Likewise, these 25 genes stratified HLH patients from healthy controls (**Figure 5F** and **Supplementary Figure S5B** and **S5D**). The PCA indicated that some of these DEGs (e.g., *AZU1, CEACAM8, CTSG, DEFA4, ELANE, LCN2, OLFM4, and MMP8*) as more associated with COVID-19_ICU than with COVID-19_nonICU.

To address whether these 25 DEGs strongly associated with COVID-19_ICU reflect only a specific similarity between COVID-19 and HLH, or if they are also linked to other acute inflammatory states, we investigated the differential expression of these molecules in other inflammatory syndromes and certain infectious diseases. We included additional inflammatory cohorts (GSE178388 [MIS-C] [34] and GSE73461 [KD] [35]) and different respiratory infections (GSE161731 [seasonal coronavirus other than SARS-CoV-2, influenza, bacterial pneumonia] [33]) (**Figure 6a**). Hierarchical cluster analysis showed the existence of a group of neutrophil-associated DEGs (e.g., *DEFA4, AZU1, ELANE, CTSG, CEACAM8, IL1R1, ARG1, LCN2, OLFM4, MMP8, CD177*, and *MCEMP1*) more consistently up-regulated across all cohorts included in this comparative analysis (**Figure 6B** and **Supplementary Table S13**). While patients with MIS-C, influenza and seasonal coronavirus showed a similar dysregulation pattern in just a few of this cluster of DEGs, patients with KD and with bacterial pneumonia exhibited a similar up-regulation pattern compared to COVID-19_ICU and HLH. Taken together, these data indicate that these DEGs reflect a more generalized inflammatory state rather than being specific to COVID-19 or HLH.

**Figure 6:**
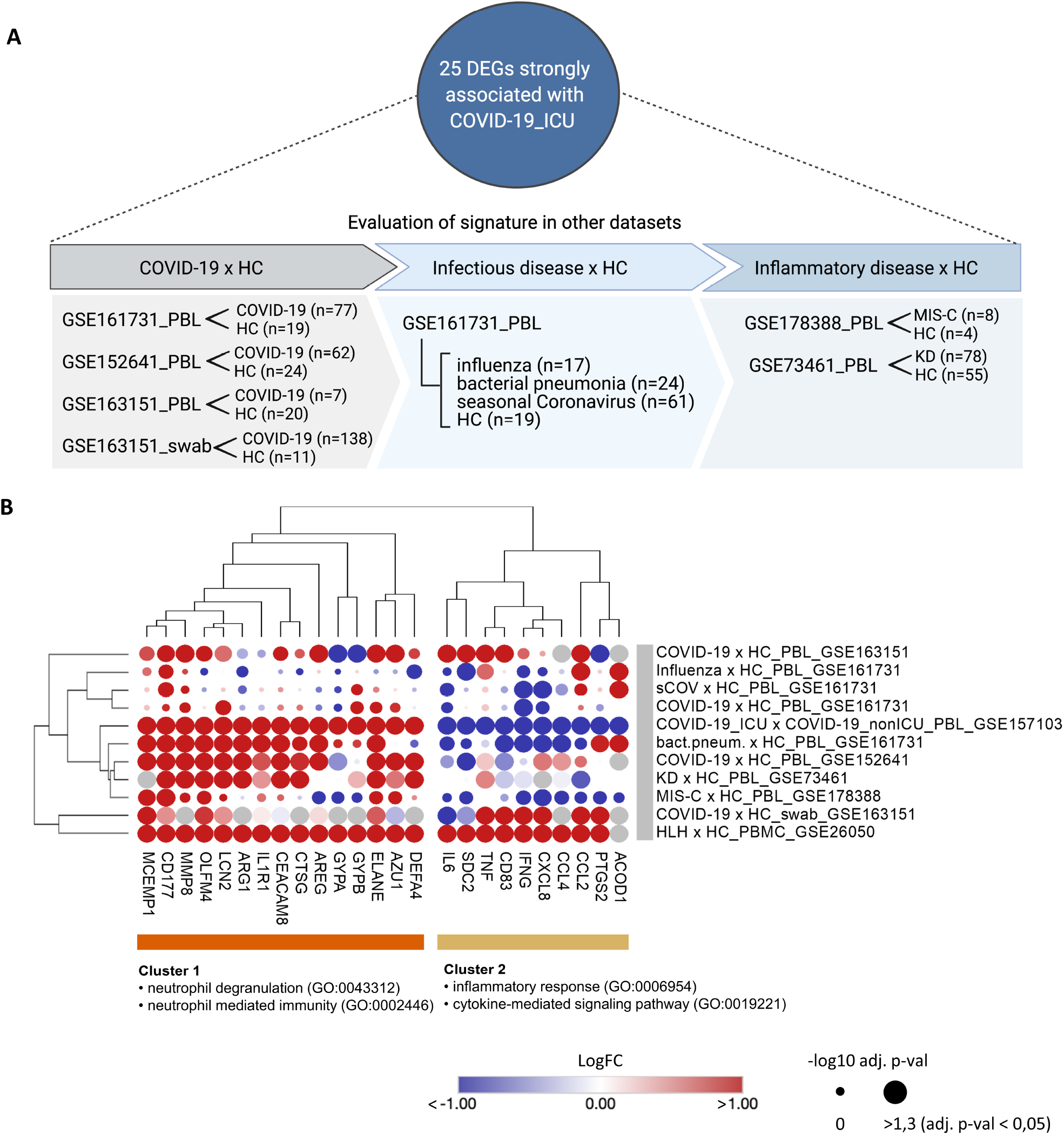
Severe COVID-19 shares a common neutrophil activation signature with other acute inflammatory states. (**A**) Schematic overview of the additional datasets included to evaluate the modulation of the 25 DEGs strongly associated with COVID-19_ICU. (**B**) Bubble heatmap showing the hierarchical clustering based on one minus spearman rank correlation of the expression pattern of these 25 DEGs across different datasets. Cluster 1 comprises genes associated with neutrophil degranulation and neutrophil-mediated immunity enriched terms, while cluster 2 includes genes enriched in inflammatory response, and cytokine-mediated signaling pathway gene ontology (GO) categories.

### Multi-layered transcriptomic analysis associates neutrophil activation signature with COVID-19 severity

Since scRNAseq allows comparison of the transcriptomes of individual cells, we next sought to investigate the distribution pattern of these 25 genes associated with COVID-19 severity. We analyzed the scRNAseq dataset (EGAS00001004571) reported by Schulte-Schrepping et al. [29] (schematic overview of study group **Figure 7A**) and found that 21 of the 25 genes associated with COVID-19 severity and HLH development are DEGs among the top 2,000 variable genes in the COVID-19 cohort compared to controls (**Figure 7B** and **Supplementary Figure S6A** and **S6B**). These 21 genes exhibited cell-type-specific expression patterns. For instance, CCL4 (a chemoattractant and stimulator of T-cell immune responses [84,85]) was mainly produced by CD8+ T and NK cells, and CD83 (B, T and dendritic cell activation marker [86,87]) by B cells and monocytes. CXCL8 was mostly present in monocytes and low-density neutrophils/granulocytes (LDG; also frequently reported as immature neutrophils [88–90]), which are neutrophils remaining in the PBMC fraction after density gradient separation. Among these 21 genes, 11 genes (among them also the 8 genes described above) were differentially expressed when comparing patients with mild and severe COVID-19 (**Figure 7C**).

**Figure 7:**
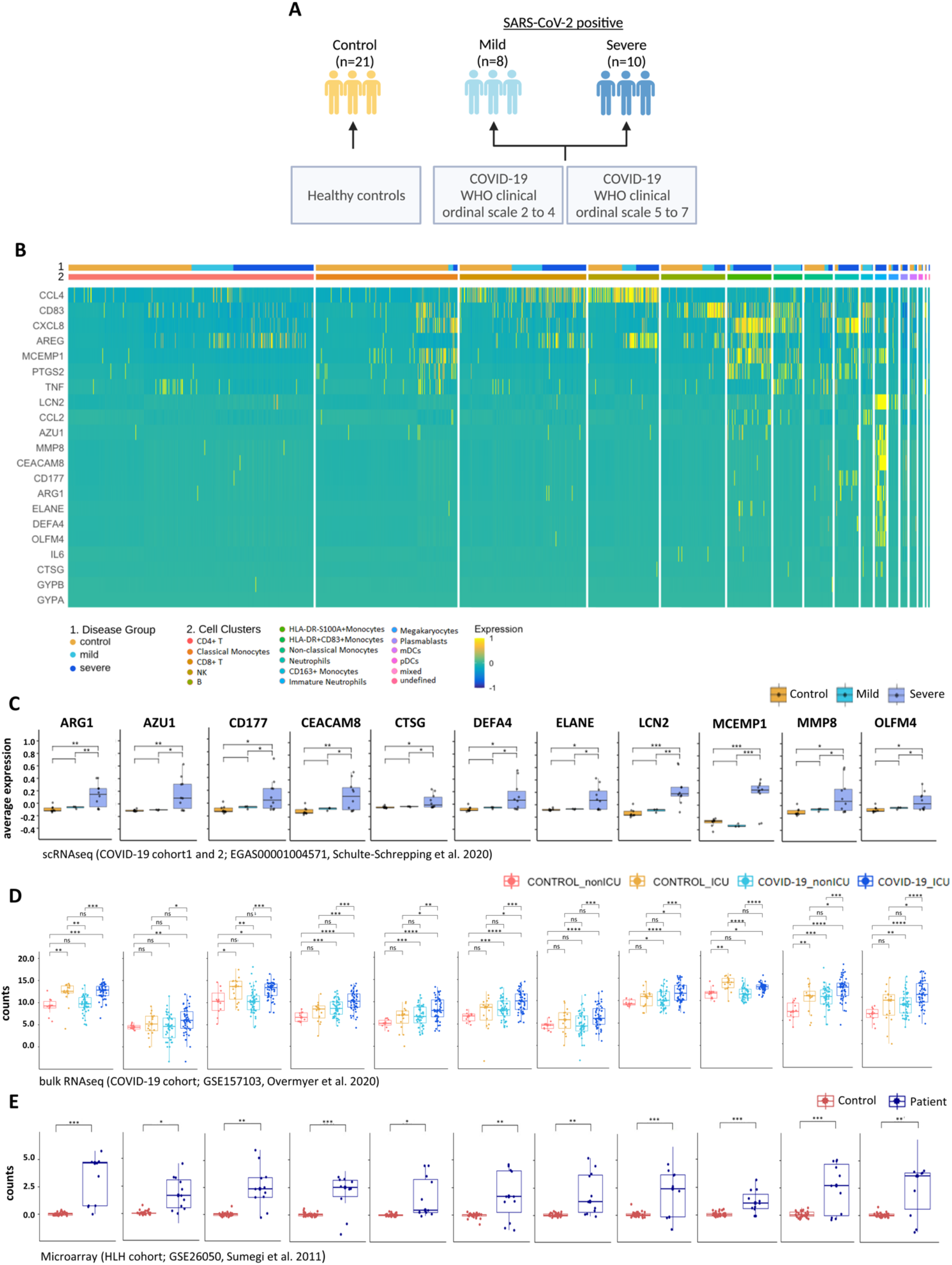
Multi-layered transcriptomic analysis associates neutrophil activation signature with COVID-19 severity. (**A**) Schematic overview of sample cohort and classification of scRNAseq dataset obtained by Schulte-Schrepping et al. [29] and used for the following analysis. **(B)** Heatmap showing scRNAseq expression of differentially expressed genes (DEGs) associated with disease severity. Cells and cohorts (controls, mild and severe COVID-19) are indicated by different colors in the legends. **(C)** Box plots of scRNAseq expression demonstrating that 11 from the 21 genes identified in **B** are up-regulated when comparing severe and mild COVID-19 patients. **(D)** Box plots of the 11 transcripts stratifying COVID-19_ICU patients from COVID-19_nonICU patients obtained from the bulk RNAseq dataset from Overmyer et al. [26]. Significant differences between groups are indicated by asterisks (* p ≤ 0.05, ** p ≤ 0.01 and *** p ≤ 0.001). (E) Box plots of microarray data illustrating that the disease severity association of COVID-19 detected by scRNAseq corresponds to the expression differences between HLH patients and controls. Significant differences between groups are indicated by asterisks (* p ≤ 0.05, ** p ≤ 0.01 and *** p ≤ 0.001).

Of note, these 11 genes encode proteins that are crucial for several pathways involved in neutrophil-mediated immunity, and are associated with diseases that increase the risk of severe COVID-19 [91,92] such as chronic obstructive pulmonary disease (COPD) [93,94] and ulcerative colitis [95,96] (**Supplementary Figure S6C** and **Supplementary Table S14**). These 11 genes are also significantly different between COVID-19_ICU and COVID-19_nonICU (**Figure 7D**) in the bulk RNAseq dataset (GSE157103, Overmyer et al. 2020 [26]), indicating that these genes are consistently associated with COVID-19 severity across different patient cohorts. Moreover, these 11 genes were differentially expressed in HLH patients compared to healthy controls (**Figure 7E**).

We used random forest method [57] to rank the importance of these 11 genes based on their ability to discriminate between COVID-19_ICU and COVID-19_nonICU in order to evaluate the association of these genes with COVID-19 severity. This approach showed an error rate (out of bag or OOB) of 27,03% and an area under the Receiver Operating Characteristic (ROC) curve of 82,4% for both groups (**Figure 8A** and **8B**). Follow-up analysis indicated that ARG1 was the most significant predictor for ICU admission followed by CD177, MCEMP1, LCN2, AZU1, OLFM4, MMP8, ELANE, CTSG, DEFA4, CEACAM8 based on the number of the nodes, gini-decrease, and average depth criteria for measuring gene importance (**Figure 8C and 8D**). ARG1 exhibited the most relevant interactions with the other genes according to the mean minimal depth criterion, mostly interacting with CD177, AZU1, MCEMP1, and LCN2 (**Figure 8E**).

**Figure 8:**
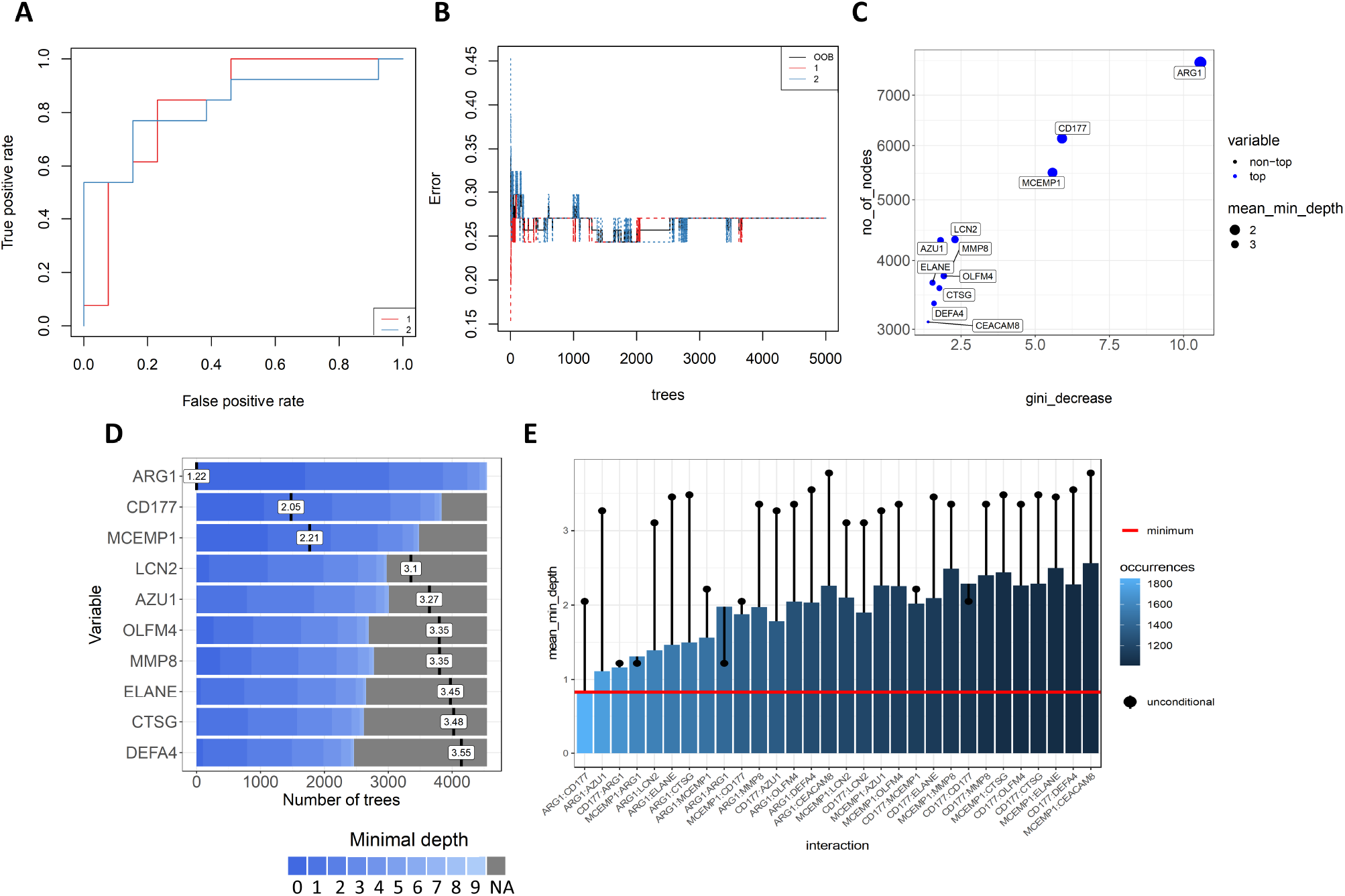
Random Forest prediction analysis suggests potential biomarker for severe COVID-19. **(A)** Receiver operating characteristics (ROC) curve of 11 genes from COVID-19_ICU compared to COVID-19_nonICU patients with an area under the curve (AUC) of 82,4% for both groups. 1=COVID-19_nonICU; 2=COVID-19_ICU. **(B)** Stable Curve showing number of trees and error rate (out of bag or OOB) with medium of 27,03%. 1=COVID-19_nonICU; 2=COVID-19_ICU. **(C)** Variable importance scores plot based on gini_decrease and number(no)_of_nodes for each variable showing which variables are more likely to be essential in the random forest’s prediction. **(D)** Ranking of the top 10 variables according to mean minimal depth (vertical bar with the mean value in it) calculated using trees. The blue color gradient reveals the min and max minimal depth for each variable. The range of the x-axis is from zero to the maximum number of trees for the feature. **(E)** Mean minimal depth variable interaction plot showing most frequent occurring interactions between the variables on the left side with light blue color, and least frequent occurring interactions on the right side of the graph with dark blue color. The red horizontal line indicates the smallest mean minimum depth and the black lollipop represent the unconditional mean minimal depth of a variable.

## 4. Discussion

Our meta-analysis integrates and unravels the consistency of several important individual studies and datasets that validate the transcriptome data also at the protein level in COVID-19 patients [26,29,97]. In agreement with the recent observation that neutrophil hyperactivation plays a key role in the severity of COVID-19 [98–101], our study indicates that severe COVID-19 disease shares a common neutrophil activation signature with other different acute inflammatory conditions such as HLH [102,103], KD [104–106], and bacterial pneumonia [107]. Our data are in agreement with the dual role of neutrophils in providing essential antimicrobial functions, as well as initiating tissue injury caused by immune dysregulation [108,109]. The genes associated with COVID-19 severity are up-regulated across different leukocyte subpopulations such as lymphoid (NK, T and B cells) and myeloid (monocytes, dendritic cells and LDGs) cells. They form a systemic and interconnected network of cell-type-specific expression pattern and signaling networks that may contribute to the clinical similarities between COVID-19 and other inflammatory conditions. Thus, our analysis identified new candidate biomarkers and novel putative molecular pathways that could lead to novel therapeutic interventions for COVID-19.

Our work expands the efforts of others [110–114] and our group [21] to identify networks and pathways involved in the pathogenesis of severe COVID-19. In accordance with our findings, it has recently been demonstrated that neutrophils accumulate in inflamed tissues of COVID-19 patients as a consequence of T-cell driven pro-inflammatory cytokine and chemokine release, which does not return to a homeostatic level due to an ineffective T cell cytotoxic response [102,103]. Moreover, our multi-layered transcriptomics approach is in agreement with the computational model developed by Ding et al. [103], which is based on a network-informed analysis of the interaction of SARS-CoV-2 and HLH related genes. This model postulates that neutrophil degranulation/NETs cause endothelial damage, and consequently, thrombotic complications of COVID-19. Ding’s and our interpretation is supported by experimental evidence [98–100,115] for neutrophil hyperactivation and its association with the severity of COVID-19, as recently reviewed by Ackermann et al. [101]. As we were able to demonstrate by the multi-omics association between leukocyte and plasma molecules, recently published flow cytometry and proteomic data indicate a systemic and integrated network of molecules associated with neutrophil growth, activation, and mobilization leading to neutrophil dysregulation in severe COVID-19 [99,100]. These results support the concept that the pathophysiology of HLH does not only involve T cell, NK cell and macrophage dysregulation, but also hyperactivation of neutrophils, as this is also seen in patients with KD [104–106] and bacterial pneumonia [107].

Among the common neutrophil activation signature that is shared by COVID-19 patients and those with other acute inflammatory states, 11 genes (*ARG1, AZU1, CD177, CEACAM8, CTSG, DEFA4, ELANE, LCN2, MCEMP1, MMP8*, and *OLFM4*) commonly dysregulated in COVID-19 and HLH specifically stratified COVID-19_ICU from COVID-19_nonICU patients. They encode proteins involved in neutrophil degranulation and contribute to the development of comorbidities that increase the risk of progressing to severe COVID-19 [91,92]. Random forest model ranking indicated that these genes accurately distinguish COVID-19_nonICU from COVID-19_ICU patients. For instance, this machine learning approach ranked ARG1 and its interaction with other molecules (CD177, AZU1, MCEMP1 and LCN2) as an important predictor for ICU admission, supporting the role of these molecules as biomarkers for hyperinflammatory conditions including those associated with severe COVID-19 [97,116].

It is important to mention that our work requires future mechanistic investigation. However, in support of our findings, several of the dysregulated molecules shared by COVID-19 and other acute inflammatory states have been successfully investigated for the treatment of SARS-CoV-2 infection. For instance, inhibition of the CCR5-CCL4 axis by Leronlima (anti-CCR5 monoclonal antibody) [117], or blockade of cytokine signaling by Tocilizumab (anti-IL-6R) [118], Adalimumab (anti-TNF) [119], or Anakinra (anti-IL1R) [120] have been shown to ameliorate, in some cases, severe COVID-19. Furthermore, Ruxolitinib, a JAK1/JAK2 inhibitor acting downstream of JAK-dependent chemokines/cytokines such as IFN-γ, IL-1β, IL-6, TNF, G-CSF, CXCL9, and CXCL10 [102,121] has shown promising results in treating COVID-19 [122]. Of note, several approaches targeting neutrophils to treat SARS-CoV-2 complications have entered clinical trials, including the disruption of signaling via CXCR2, IL-8, IL-17A, or the use of phosphodiesterase (PDE) inhibitors [123]. Moreover, in agreement with our data, inhibition of neutrophil-derived anti-microbial proteins are being actively investigated in clinical trials by exploring the mechanistic and clinical effects of Alvelestat, an oral neutrophil elastase inhibitor (COVID-19 Study of Safety and Tolerability of Alvelestat, ClinicalTrials.gov). Meanwhile other proteases (AZU1 and CTSG) and the inhibition of NET formation have been suggested to alleviate SARS-CoV-2 symptoms [97,101,124].

In conclusion, our comprehensive multi-layered transcriptomic and cross-tissue analysis indicates systemic communalities among severe COVID-19 and other acute inflammatory states. This work suggests that an interconnected cytokine/chemokine profile that hyper stimulates and systemically attracts adaptive and innate immune cells, culminating in the hyperactivation of neutrophils. Altogether, these data indicate that both numeric and dysfunctional changes of neutrophils [125,126] are involved in COVID-19 outcomes, i.e., high levels of circulating activated neutrophils [125,127]. Thus, our work suggests common molecular pathways between severe COVID-19 and other acute inflammatory states that can be exploited for therapeutic intervention.

## Supplementary Materials

All supplemental figures and legends are provided in a separate document file. Figure S1: Selection of common differentially expressed genes (DEGs) across different COVID-19 datasets and the HLH dataset; Figure S2: Molecular relationships between COVID-19 and fHLH; Figure S3: Bivariate correlation of cytokine/chemotaxis and neutrophil-mediated immunity genes; Figure S4: Significant immune pathways and dysregulated molecules obtained from proteomics analyses of plasma from severe COVID-19 when compared to mild disease; Figure S5: Contribution of genes and individuals to stratification of severe COVID-19 and HLH; Figure S6: Cell cluster profile of scRNA seq analysis and enrichment of 11 genes significant for severe COVID-19.

Supplemental Tables S1-S14 are provided in a separate document file. Table S1: Up- and Down-regulated DEGs of all datasets used for defining common DEGs; Table S2: List of common DEGs; Table S3: Genelist overlaps; Table S4: Gene Ontology Biological Process (BP) enriched terms; Table S5: Gene Ontology Cellular Components (CC) enriched terms; Table S6: Cytokine and Chemotaxis genes; Table S7: Neutrophil-mediated immunity genes; Table S8: Pleiotropic genes; Table S9: DEGs between COVID-19_ICU and COVID-19_nonICU; Table S10: 25 Genes in proteomic/protein expression datasets; Table S11: Significant plasma proteins comparing COVID-19_ICU versus COVID-19_nonICU patients; Table S12: Enriched BP GO terms of 158 significant plasma proteins; Table S13: 25 DEGs in additional datasets; Table S14: Enriched Elsevier Pathway collection terms of 11 genes significant for severe COVID-19

## Author Contributions

LFS, and OCM wrote the manuscript, conceived the study design, bioinformatics analyses and performed scientific insights; AHCM, GCB, CASP, DLMF, PPF, DRP, ISF, RCS, GJM, AERO, and IJ, co-wrote the manuscript and performed bioinformatics analyses; KAO, JJC, JB, AJ, JSS, TU and JLS generated transcriptome and proteome data; JPSP, GCM, JAMB, NOSC, HDO, VLGC, ACN, KAO, JJC, JB, AJ, JSS, TU, JLS, HIN, IJ and OCM revised and edited the final manuscript and provided scientific input; OCM supervised the project.

## Funding and acknowledgements

We acknowledge the Latin American Society of Immunodeficiencies (LASID) for providing the research funding of LFS (LASID Fellowship award 2020), and the São Paulo Research Foundation (FAPESP grants 2018/18886-9, 2020/01688-0, and 2020/07069-0 to OCM) for financial support. Computational analysis was supported by FAPESP and partially by the grants from Ontario Research Fund (#34876), Natural Sciences Research Council (NSERC #203475), Canada Foundation for Innovation (CFI #29272, #225404, #33536), and IBM granted to IJ, the National Institutes of Health (NHLBI) through award HL130704 and HL160661 granted to AJ, as well as the NIH P41 GM108538 granted to KAO and JJC. This study was financed in part by the coordination for the improvement of higher education personnel – Brazil (CAPES) – finance code 001. We thank Prof. Luis Carlos de Souza Ferreira for discussion and suggestions to develop this project.

## Data Availability Statement

This paper analyzes existing, publicly available data. The accession numbers for the datasets are listed in the key resources table.

All original codes used for data analysis have been deposited at github (https://github.com/lschimke/COVID19-and-HLH-paper) and are publicly available as of the date of publication. R packages are listed in the key resources table.

Any additional information required to reanalyze the data reported in this paper is available from the lead contact upon request.

## References

1. Mehta, P.; McAuley, D.F.; Brown, M.; Sanchez, E.; Tattersall, R.S.; Manson, J.J. COVID-19: consider cytokine storm syndromes and immunosuppression. Lancet 2020, 395, 1033–1034.

2. Chen, L.Y.C.; Quach, T.T.T. COVID-19 cytokine storm syndrome: a threshold concept. 2021, doi:10.1016/j.blre.2020.100707.

3. Huang, C.; Wang, Y.; Li, X.; Ren, L.; Zhao, J.; Hu, Y.; Zhang, L.; Fan, G.; Xu, J.; Gu, X.; et al. Clinical features of patients infected with 2019 novel coronavirus in Wuhan, China. Lancet 2020, 395, 497–506, doi:10.1016/S0140-6736(20)30183-5.

4. England, J.T.; Abdulla, A.; Biggs, C.M.; Lee, A.Y.Y.; Hay, K.A.; Hoiland, R.L.; Wellington, C.L.; Sekhon, M.; Jamal, S.; Shojania, K.; et al. Since January 2020 Elsevier has created a COVID-19 resource centre with free information in English and Mandarin on the novel coronavirus COVID-19. The COVID-19 resource centre is hosted on Elsevier Connect, the company ’s public news and information. 2020.

5. Ørskov, S.; Frost Nielsen, B.; Føns, S.; Sneppen, K.; Simonsen, L. The COVID-19 pandemic: Key considerations for the epidemic and its control. APMIS 2021, doi:10.1111/apm.13141.

6. Pandit, B.; Bhattacharjee, S.; Bhattacharjee, B. Association of clade-G SARS-CoV-2 viruses and age with increased mortality rates across 57 countries and India. Infect. Genet. Evol. 2021, 90, doi:10.1016/j.meegid.2021.104734.

7. Garcia-Beltran, W.F.; Lam, E.C.; St. Denis, K.; Nitido, A.D.; Garcia, Z.H.; Hauser, B.M.; Feldman, J.; Pavlovic, M.N.; Gregory, D.J.; Poznansky, M.C.; et al. Multiple SARS-CoV-2 variants escape neutralization by vaccine-induced humoral immunity. Cell 2021, 184, 2372–2383.e9, doi:10.1016/j.cell.2021.03.013.

8. Kalfaoglu, B.; Almeida-Santos, J.; Tye, C.A.; Satou, Y.; Ono, M. T-Cell Hyperactivation and Paralysis in Severe COVID-19 Infection Revealed by Single-Cell Analysis. Front. Immunol. 2020, 11, 2605, doi:10.3389/FIMMU.2020.589380/BIBTEX.

9. Wang, E.Y.; Mao, T.; Klein, J.; Dai, Y.; Huck, J.D.; Jaycox, J.R.; Liu, F.; Zhou, T.; Israelow, B.; Wong, P.; et al. Diverse functional autoantibodies in patients with COVID-19. Nat. 2021 5957866 2021, 595, 283–288, doi:10.1038/s41586-021-03631-y.

10. Cabral-Marques, O.; Halpert, G.; Schimke, L.F.; Ostrinski, Y.; Zyskind, I.; Lattin, M.T.; Tran, F.; Schreiber, S.; Marques, A.H.C.; Filgueiras, I.S.; et al. The relationship between autoantibodies targeting GPCRs and the renin-angiotensin system associates with COVID-19 severity. medRxiv 2021, 2021.08.24.21262385, doi:10.1101/2021.08.24.21262385.

11. Mathew, D.; Giles, J.R.; Baxter, A.E.; Oldridge, D.A.; Greenplate, A.R.; Wu, J.E.; Alanio, C.; Kuri-Cervantes, L.; Pampena, M.B.; D’Andrea, K.; et al. Deep immune profiling of COVID-19 patients reveals distinct immunotypes with therapeutic implications. Science (80-.). 2020, 369, doi:10.1126/SCIENCE.ABC8511.

12. Laing, A.G.; Lorenc, A.; del Molino del Barrio, I.; Das, A.; Fish, M.; Monin, L.; Muñoz-Ruiz, M.; McKenzie, D.R.; Hayday, T.S.; Francos-Quijorna, I.; et al. A dynamic COVID-19 immune signature includes associations with poor prognosis. Nat. Med. 2020, 26, 1623–1635, doi:10.1038/s41591-020-1038-6.

13. Azkur, A.K.; Akdis, M.; Azkur, D.; Sokolowska, M.; van de Veen, W.; Brüggen, M.C.; O’Mahony, L.; Gao, Y.; Nadeau, K.; Akdis, C.A. Immune response to SARS-CoV-2 and mechanisms of immunopathological changes in COVID-19. Allergy Eur. J. Allergy Clin. Immunol. 2020, 75, 1564–1581.

14. Koutsakos, M.; Rowntree, L.C.; Hensen, L.; Chua, B.Y.; van de Sandt, C.E.; Habel, J.R.; Zhang, W.; Jia, X.; Kedzierski, L.; Ashhurst, T.M.; et al. Integrated immune dynamics define correlates of COVID-19 severity and antibody responses. Cell Reports Med. 2021, 2, doi:10.1016/j.xcrm.2021.100208.

15. Webb, B.J.; Peltan, I.D.; Jensen, P.; Hoda, D.; Hunter, B.; Silver, A.; Starr, N.; Buckel, W.; Grisel, N.; Hummel, E.; et al. Clinical criteria for COVID-19-associated hyperinflammatory syndrome: a cohort study. Lancet Rheumatol. 2020, 2, e754–e763, doi:10.1016/S2665-9913(20)30343-X.

16. Fajgenbaum, D.C.; June, C.H. Cytokine Storm. N. Engl. J. Med. 2020, 383, 2255–2273, doi:10.1056/nejmra2026131.

17. Yongzhi, X. COVID-19-associated cytokine storm syndrome and diagnostic principles: an old and new Issue. Emerg. Microbes Infect. 2021, 10, 266–276.

18. George, M. Hemophagocytic lymphohistiocytosis: review of etiologies and management. J. Blood Med. 2014, 69, doi:10.2147/jbm.s46255.

19. Soy, M.; Atagündüz, P.; Atagündüz, I.; Sucak, G.T. Hemophagocytic lymphohistiocytosis: a review inspired by the COVID-19 pandemic. Rheumatol. Int. 2021, 41, 7–18, doi:10.1007/s00296-020-04636-y.

20. Lorenz, G.; Moog, P.; Bachmann, Q.; La Rosée, P.; Schneider, H.; Schlegl, M.; Spinner, C.; Heemann, U.; Schmid, R.M.; Algül, H.; et al. Title: Cytokine release syndrome is not usually caused by secondary hemophagocytic lymphohistiocytosis in a cohort of 19 critically ill COVID-19 patients. Sci. Rep. 2020, 10, 18277, doi:10.1038/s41598-020-75260-w.

21. Freire, P.P.; Marques, A.H.C.; Baiocchi, G.C.; Schimke, L.F.; Fonseca, D.L.M.; Salgado, R.C.; Filgueiras, I.S.; Napoleao, S.M.S.; Plaça, D.R.; Akashi, K.T.; et al. The relationship between cytokine and neutrophil gene network distinguishes SARS-CoV-2–infected patients by sex and age. JCI Insight 2021, 6, doi:10.1172/jci.insight.147535.

22. Alqutami, F.; Senok, A.; Hachim, M. COVID-19 Transcriptomic Atlas: A Comprehensive Analysis of COVID-19 Related Transcriptomics Datasets. Front. Genet. 2021, 12, doi:10.3389/FGENE.2021.755222.

23. Delorey, T.M.; Ziegler, C.G.K.; Heimberg, G.; Normand, R.; Yang, Y.; Segerstolpe, Å.; Abbondanza, D.; Fleming, S.J.; Subramanian, A.; Montoro, D.T.; et al. COVID-19 tissue atlases reveal SARS-CoV-2 pathology and cellular targets. Nat. 2021 5957865 2021, 595, 107–113, doi:10.1038/s41586-021-03570-8.

24. Daamen, A.R.; Bachali, P.; Owen, K.A.; Kingsmore, K.M.; Hubbard, E.L.; Labonte, A.C.; Robl, R.; Shrotri, S.; Grammer, A.C.; Lipsky, P.E. Comprehensive transcriptomic analysis of COVID-19 blood, lung, and airway. Sci. Reports 2021 111 2021, 11, 1–19, doi:10.1038/s41598-021-86002-x.

25. Arunachalam, P.S.; Wimmers, F.; Mok, C.K.P.; Perera, R.A.P.M.; Scott, M.; Hagan, T.; Sigal, N.; Feng, Y.; Bristow, L.; Tsang, O.T.Y.; et al. Systems biological assessment of immunity to mild versus severe COVID-19 infection in humans. Science (80-.). 2020, 369, 1210–1220, doi:10.1126/SCIENCE.ABC6261.

26. Overmyer, K.A.; Shishkova, E.; Miller, I.J.; Balnis, J.; Bernstein, M.N.; Peters-Clarke, T.M.; Meyer, J.G.; Quan, Q.; Muehlbauer, L.K.; Trujillo, E.A.; et al. Large-Scale Multi-omic Analysis of COVID-19 Severity. Cell Syst. 2020, doi:10.1016/j.cels.2020.10.003.

27. Lieberman, N.A.P.; Peddu, V.; Xie, H.; Shrestha, L.; Huang, M.-L.; Mears, M.C.; Cajimat, M.N.; Bente, D.A.; Shi, P.-Y.; Bovier, F.; et al. In vivo antiviral host transcriptional response to SARS-CoV-2 by viral load, sex, and age. PLoS Biol. 2020, 18, doi:10.1371/JOURNAL.PBIO.3000849.

28. Mick, E.; Kamm, J.; Pisco, A.O.; Ratnasiri, K.; Babik, J.M.; Calfee, C.S.; Castaneda, G.; DeRisi, J.L.; Detweiler, A.M.; Hao, S.; et al. Upper airway gene expression differentiates COVID-19 from other acute respiratory illnesses and reveals suppression of innate immune responses by SARS-CoV-2. medRxiv Prepr. Serv. Heal. Sci. 2020, doi:10.1101/2020.05.18.20105171.

29. Schulte-Schrepping, J.; Reusch, N.; Paclik, D.; Baßler, K.; Schlickeiser, S.; Zhang, B.; Krämer, B.; Krammer, T.; Brumhard, S.; Bonaguro, L.; et al. Severe COVID-19 Is Marked by a Dysregulated Myeloid Cell Compartment. Cell 2020, doi:10.1016/j.cell.2020.08.001.

30. Sumegi, J.; Barnes, M.G.; Nestheide, S. V.; Molleran-Lee, S.; Villanueva, J.; Zhang, K.; Risma, K.A.; Grom, A.A.; Filipovich, A.H. Gene expression profiling of peripheral blood mononuclear cells from children with active hemophagocytic lymphohistiocytosis. Blood 2011, 117, doi:10.1182/blood-2010-08-300046.

31. Ng, D.; Granados, A.; Santos, Y.; Servellita, V.; Goldgof, G.; Meydan, C.; Sotomayor-Gonzalez, A.; Levine, A.; Balcerek, J.; Han, L.; et al. A diagnostic host response biosignature for COVID-19 from RNA profiling of nasal swabs and blood. Sci. Adv. 2021, 7, doi:10.1126/SCIADV.ABE5984.

32. Thair, S.A.; He, Y.D.; Hasin-Brumshtein, Y.; Sakaram, S.; Pandya, R.; Toh, J.; Rawling, D.; Remmel, M.; Coyle, S.; Dalekos, G.N.; et al. Transcriptomic similarities and differences in host response between SARS-CoV-2 and other viral infections. iScience 2021, 24,doi:10.1016/j.isci.2020.101947.

33. McClain, M.T.; Constantine, F.J.; Henao, R.; Liu, Y.; Tsalik, E.L.; Burke, T.W.; Steinbrink, J.M.; Petzold, E.; Nicholson, B.P.; Rolfe, R.; et al. Dysregulated transcriptional responses to SARS-CoV-2 in the periphery. Nat. Commun. 2021 121 2021, 12, 1–8, doi:10.1038/s41467-021-21289-y.

34. Beckmann, N.D.; Comella, P.H.; Cheng, E.; Lepow, L.; Beckmann, A.G.; Tyler, S.R.; Mouskas, K.; Simons, N.W.; Hoffman, G.E.; Francoeur, N.J.; et al. Downregulation of exhausted cytotoxic T cells in gene expression networks of multisystem inflammatory syndrome in children. Nat. Commun. 2021 121 2021, 12, 1–15, doi:10.1038/s41467-021-24981-1.

35. Wright, V.J.; Herberg, J.A.; Kaforou, M.; Shimizu, C.; Eleftherohorinou, H.; Shailes, H.; Barendregt, A.M.; Menikou, S.; Gormley, S.; Berk, M.; et al. Diagnosis of Kawasaki Disease Using a Minimal Whole-Blood Gene Expression Signature. JAMA Pediatr. 2018, 172, doi:10.1001/JAMAPEDIATRICS.2018.2293.

36. Clough, E.; Barrett, T. The Gene Expression Omnibus database. In Methods in Molecular Biology; Humana Press Inc., 2016; Vol. 1418, pp. 93–110.

37. Athar, A.; Füllgrabe, A.; George, N.; Iqbal, H.; Huerta, L.; Ali, A.; Snow, C.; Fonseca, N.A.; Petryszak, R.; Papatheodorou, I.; et al. ArrayExpress update - From bulk to single-cell expression data. Nucleic Acids Res. 2019, 47, D711–D715, doi:10.1093/nar/gky964.

38. Sanchis, P.; Lavignolle, R.; Abbate, M.; Lage-Vickers, S.; Vazquez, E.; Cotignola, J.; Bizzotto, J.; Gueron, G. Analysis workflow of publicly available RNA-sequencing datasets. STAR Protoc. 2021, 2, 100478, doi:10.1016/J.XPRO.2021.100478.

39. Humaidan, P.; Polyzos, N.P. (Meta)analyze this: Systematic reviews might lose credibility. Nat. Med. 2012 189 2012, 18, 1321–1321, doi:10.1038/nm0912-1321.

40. Zhou, G.; Soufan, O.; Ewald, J.; Hancock, R.E.W.; Basu, N.; Xia, J. NetworkAnalyst 3.0: A visual analytics platform for comprehensive gene expression profiling and meta-analysis. Nucleic Acids Res. 2019, 47, W234–W241, doi:10.1093/nar/gkz240.

41. Law, C.W.; Chen, Y.; Shi, W.; Smyth, G.K. Voom: Precision weights unlock linear model analysis tools for RNA-seq read counts. Genome Biol. 2014, 15, R29, doi:10.1186/gb-2014-15-2-r29.

42. Bardou, P.; Mariette, J.; Escudié, F.; Djemiel, C.; Klopp, C. Jvenn: An interactive Venn diagram viewer. BMC Bioinformatics 2014, 15, doi:10.1186/1471-2105-15-293.

43. Krzywinski, M.; Schein, J.; Birol, I.; Connors, J.; Gascoyne, R.; Horsman, D.; Jones, S.J.; Marra, M.A. Circos: An information aesthetic for comparative genomics. Genome Res. 2009, 19, 1639–1645, doi:10.1101/gr.092759.109.

44. Hao, Y.; Hao, S.; Andersen-Nissen, E.; Mauck, W.M.; Zheng, S.; Butler, A.; Lee, M.J.; Wilk, A.J.; Darby, C.; Zagar, M.; et al. Integrated analysis of multimodal single-cell data. bioRxiv 2020, 2020.10.12.335331.

45. Stuart, T.; Butler, A.; Hoffman, P.; Hafemeister, C.; Papalexi, E.; Mauck, W.M.; Hao, Y.; Stoeckius, M.; Smibert, P.; Satija, R. Comprehensive Integration of Single-Cell Data. Cell 2019, 177, 1888–1902.e21, doi:10.1016/j.cell.2019.05.031.

46. Brown, K.R.; Otasek, D.; Ali, M.; McGuffin, M.J.; Xie, W.; Devani, B.; van Toch, I.L.; Jurisica, I. NAViGaTOR: Network Analysis, Visualization and Graphing Toronto. Bioinformatics 2009, 25, 3327–3329, doi:10.1093/BIOINFORMATICS/BTP595.

47. Kotlyar, M.; Pastrello, C.; Ahmed, Z.; Chee, J.; Varyova, Z.; Jurisica, I. IID 2021: towards context-specific protein interaction analyses by increased coverage, enhanced annotation and enrichment analysis. Nucleic Acids Res. 2022, 50, D640–D647, doi:10.1093/NAR/GKAB1034.

48. Yu, G.; Wang, L.G.; Han, Y.; He, Q.Y. ClusterProfiler: An R package for comparing biological themes among gene clusters. Omi. A J. Integr. Biol. 2012, 16, 284–287, doi:10.1089/omi.2011.0118.

49. Chen, E.Y.; Tan, C.M.; Kou, Y.; Duan, Q.; Wang, Z.; Meirelles, G. V.; Clark, N.R.; Ma’ayan, A. Enrichr: Interactive and collaborative HTML5 gene list enrichment analysis tool. BMC Bioinformatics 2013, 14, doi:10.1186/1471-2105-14-128.

50. Kuleshov, M. V.; Jones, M.R.; Rouillard, A.D.; Fernandez, N.F.; Duan, Q.; Wang, Z.; Koplev, S.; Jenkins, S.L.; Jagodnik, K.M.; Lachmann, A.; et al. Enrichr: a comprehensive gene set enrichment analysis web server 2016 update. Nucleic Acids Res. 2016, 44, W90–W97, doi:10.1093/nar/gkw377.

51. Xie, Z.; Bailey, A.; Kuleshov, M. V.; Clarke, D.J.B.; Evangelista, J.E.; Jenkins, S.L.; Lachmann, A.; Wojciechowicz, M.L.; Kropiwnicki, E.; Jagodnik, K.M.; et al. Gene Set Knowledge Discovery with Enrichr. Curr. Protoc. 2021, 1, e90, doi:10.1002/cpz1.90.

52. Starruß, J.; de Back, W.; Brusch, L.; Deutsch, A. Morpheus: a user-friendly modeling environment for multiscale and multicellular systems biology. Bioinformatics 2014, 30, 1331–2, doi:10.1093/bioinformatics/btt772.

53. Kassambara, A.; Mundt, F. Multivariate Analysis II, Practical Guide to Principal Component Methods in R; 1st, 2017th ed.; STHDA;

54. Jendoubi, T.; Strimmer, K. A whitening approach to probabilistic canonical correlation analysis for omics data integration. BMC Bioinformatics 2019, 20, 15, doi:10.1186/s12859-018-2572-9.

55. Konietschke, F.; Placzek, M.; Schaarschmidt, F.; Hothorn, L.A. nparcomp: An R Software Package for Nonparametric Multiple Comparisons and Simultaneous Confidence Intervals. J. Stat. Softw. 2015, 64, 1–17, doi:10.18637/JSS.V064.I09.

56. Burchett, W.W.; Ellis, A.R.; Harrar, S.W.; Bathke, A.C. Nonparametric Inference for Multivariate Data: The R Package npmv. J. Stat. Softw. 2017, 76, 1–18, doi:10.18637/JSS.V076.I04.

57. Liaw, A.; Wiener, M. Classification and Regression by randomForest. 2002, 2.

58. Usmani, G.N.; Woda, B.A.; Newburger, P.E. Advances in understanding the pathogenesis of HLH. Br. J. Haematol. 2013, 161, 609–622.

59. Stelzer, G.; Rosen, N.; Plaschkes, I.; Zimmerman, S.; Twik, M.; S, F.; TI, S.; R, N.; I, L.; Y, M.; et al. The GeneCards Suite: From Gene Data Mining to Disease Genome Sequence Analyses. Curr. Protoc. Bioinforma. 2016, 54, 1.30.1–1.30.33, doi:10.1002/CPBI.5.

60. Lebre, M.C.; Burwell, T.; Vieira, P.L.; Lora, J.; Coyle, A.J.; Kapsenberg, M.L.; Clausen, B.E.; De Jong, E.C. Differential expression of inflammatory chemokines by Th1- and Th2-cell promoting dendritic cells: A role for different mature dendritic cell populations in attracting appropriate effector cells to peripheral sites of inflammation. Immunol. Cell Biol. 2005, 83, 525–535, doi:10.1111/j.1440-1711.2005.01365.x.

61. Charo, I.F.; Ransohoff, R.M. The Many Roles of Chemokines and Chemokine Receptors in Inflammation. N. Engl. J. Med. 2006, 354, 610–621, doi:10.1056/nejmra052723.

62. Blandino-Rosano, M.; Perez-Arana, G.; Mellado-Gil, J.M.; Segundo, C.; Aguilar-Diosdado, M. Anti-proliferative effect of pro-inflammatory cytokines in cultured β cells is associated with extracellular signal-regulated kinase 1/2 pathway inhibition: Protective role of glucagon-like peptide −1. J. Mol. Endocrinol. 2008, 41, 35–44, doi:10.1677/JME-07-0154.

63. Zhang, J.M.; An, J. Cytokines, inflammation, and pain. Int. Anesthesiol. Clin. 2007, 45, 27–37.

64. Aratani, Y. Myeloperoxidase: Its role for host defense, inflammation, and neutrophil function. Arch. Biochem. Biophys. 2018, 640, 47–52.

65. Pohl, J.; Pereira, H.A.; Martin, N.M.; Spitznagel, J.K. Amino acid sequence of CAP37, a human neutrophil granule-derived antibacterial and monocyte-specific chemotactic glycoprotein structurally similar to neutrophil elastase. FEBS Lett. 1990, 272, 200–204, doi:10.1016/0014-5793(90)80484-Z.

66. Korkmaz, B.; Horwitz, M.S.; Jenne, D.E.; Gauthier, F. Neutrophil elastase, proteinase 3, and cathepsin G as therapeutic targets in human diseases. Pharmacol. Rev. 2010, 62, 726–759.

67. Tralau, T.; Meyer-Hoffert, U.; Schröder, J.M.; Wiedow, O. Human leukocyte elastase and cathepsin G are specific inhibitors of C5a-dependent neutrophil enzyme release and chemotaxis. Exp. Dermatol. 2004, 13, 316–325, doi:10.1111/j.0906-6705.2004.00145.x.

68. Lehrer, R.I.; Lu, W. α-Defensins in human innate immunity. Immunol. Rev. 2012, 245, 84–112.

69. Bradley, L.M.; Douglass, M.F.; Chatterjee, D.; Akira, S.; Baaten, B.J.G. Matrix metalloprotease 9 mediates neutrophil migration into the airways in response to influenza virus-induced toll-like receptor signaling. PLoS Pathog. 2012, 8, e1002641, doi:10.1371/journal.ppat.1002641.

70. Lin, W.C.; Fessler, M.B. Regulatory mechanisms of neutrophil migration from the circulation to the airspace. Cell. Mol. Life Sci. 2021, 1, 3.

71. Cabral-Marques, O.; Schimke, L.F.; de Oliveira, E.B.; El Khawanky, N.; Ramos, R.N.; Al-Ramadi, B.K.; Segundo, G.R.S.; Ochs, H.D.; Condino-Neto, A. Flow Cytometry Contributions for the Diagnosis and Immunopathological Characterization of Primary Immunodeficiency Diseases With Immune Dysregulation. Front. Immunol. 2019, 10, 2742.

72. Marsh, R.A.; Satake, N.; Biroschak, J.; Jacobs, T.; Johnson, J.; Jordan, M.B.; Bleesing, J.J.; Filipovich, A.H.; Zhang, K. STX11 mutations and clinical phenotypes of familial hemophagocytic lymphohistiocytosis in North America. Pediatr. Blood Cancer 2010, 55, 134–140, doi:10.1002/pbc.22499.

73. Wada, T.; Sakakibara, Y.; Nishimura, R.; Toma, T.; Ueno, Y.; Horita, S.; Tanaka, T.; Nishi, M.; Kato, K.; Yasumi, T.; et al. Downregulation of CD5 expression on activated CD8+ T cells in familial hemophagocytic lymphohistiocytosis with perforin gene mutations. Hum. Immunol. 2013, 74, 1579–1585, doi:10.1016/j.humimm.2013.09.001.

74. Janka, G.E. Familial and acquired hemophagocytic lymphohistiocytosis. Eur. J. Pediatr. 2007, 166, 95–109.

75. Dell’Acqua, F.; Saettini, F.; Castelli, I.; Badolato, R.; Notarangelo, L.D.; Rizzari, C. Hermansky-Pudlak syndrome type II and lethal hemophagocytic lymphohistiocytosis: Case description and review of the literature. J. Allergy Clin. Immunol. Pract. 2019, 7, 2476–2478.e5, doi:10.1016/j.jaip.2019.04.001.

76. Zhao, X.W.; Gazendam, R.P.; Drewniak, A.; Van Houdt, M.; Tool, A.T.J.; Van Hamme, J.L.; Kustiawan, I.; Meijer, A.B.; Janssen, H.; Russell, D.G.; et al. Defects in neutrophil granule mobilization and bactericidal activity in familial hemophagocytic lymphohistiocytosis type 5 (FHL-5) syndrome caused by STXBP2/Munc18-2 mutations. Blood 2013, 122, 109–111, doi:10.1182/blood-2013-03-494039.

77. D’Orlando, O.; Zhao, F.; Kasper, B.; Orinska, Z.; Müller, J.; Hermans-Borgmeyer, I.; Griffiths, G.M.; Zur Stadt, U.; Bulfone-Paus, S. Syntaxin 11 is required for NK and CD8+ T-cell cytotoxicity and neutrophil degranulation. Eur. J. Immunol. 2013, 43, 194–208, doi:10.1002/eji.201142343.

78. Zhang, Y.; Tang, W.; Zhang, H.; Niu, X.; Xu, Y.; Zhang, J.; Gao, K.; Pan, W.; Boggon, T.J.; Toomre, D.; et al. A Network of interactions enables CCM3 and STK24 to coordinate UNC13D-driven vesicle exocytosis in neutrophils. Dev. Cell 2013, 27, 215–226, doi:10.1016/j.devcel.2013.09.021.

79. Catz, S.D. The role of Rab27a in the regulation of neutrophil function. Cell. Microbiol. 2014, 16, 1301–1310, doi:10.1111/cmi.12328.

80. Rickman, J.M.; Wang, Y.; Rollett, A.D.; Harmer, M.P.; Compson, C. Data analytics using canonical correlation analysis and Monte Carlo simulation. npj Comput. Mater. 2017, 3, 26, doi:10.1038/s41524-017-0028-9.

81. Ferreira, F.; Bota, D.; Bross, A.; Mélot, C.; Vincent, J. Serial evaluation of the SOFA score to predict outcome in critically ill patients. JAMA 2001, 286, 1754–1758, doi:10.1001/JAMA.286.14.1754.

82. Lever, J.; Krzywinski, M.; Altman, N. Points of Significance: Principal component analysis. Nat. Methods 2017, 14, 641–642.

83. Ringnér, M. What is principal component analysis? Nat. Biotechnol. 2008, 26, 303–304.

84. Moser, B. T-cell memory: the importance of chemokine-mediated cell attraction. Curr. Biol. 2006, 16.

85. Castellino, F.; Huang, A.Y.; Altan-Bonnet, G.; Stoll, S.; Scheinecker, C.; Germain, R.N. Chemokines enhance immunity by guiding naive CD8+ T cells to sites of CD4+ T cell-dendritic cell interaction. Nature 2006, 440, 890–895, doi:10.1038/nature04651.

86. Pinho, M.P.; Migliori, I.K.; Flatow, E.A.; Barbuto, J.A.M. Dendritic cell membrane CD83 enhances immune responses by boosting intracellular calcium release in T lymphocytes. J. Leukoc. Biol. 2014, 95, 755–762, doi:10.1189/jlb.0413239.

87. Krzyzak, L.; Seitz, C.; Urbat, A.; Hutzler, S.; Ostalecki, C.; Gläsner, J.; Hiergeist, A.; Gessner, A.; Winkler, T.H.; Steinkasserer, A.; et al. CD83 Modulates B Cell Activation and Germinal Center Responses. J. Immunol. 2016, 196, 3581–3594, doi:10.4049/jim-munol.1502163.

88. Silvestre-Roig, C.; Fridlender, Z.G.; Glogauer, M.; Scapini, P. Neutrophil Diversity in Health and Disease. Trends Immunol. 2019, 40, 565–583.

89. Ostendorf, L.; Mothes, R.; van Koppen, S.; Lindquist, R.L.; Bellmann-Strobl, J.; Asseyer, S.; Ruprecht, K.; Alexander, T.; Niesner, R.A.; Hauser, A.E.; et al. Low-Density Granulocytes Are a Novel Immunopathological Feature in Both Multiple Sclerosis and Neuromyelitis Optica Spectrum Disorder. Front. Immunol. 2019, 10, 2725, doi:10.3389/fimmu.2019.02725.

90. Carmona-Rivera, C.; Kaplan, M.J. Low-density granulocytes: A distinct class of neutrophils in systemic autoimmunity. Semin. Immunopathol. 2013, 35, 455–463.

91. Singh, A.K.; Jena, A.; Kumar-M, P.; Sharma, V.; Sebastian, S. Risk and outcomes of coronavirus disease in patients with inflammatory bowel disease: A systematic review and meta-analysis. United Eur. Gastroenterol. J. 2021, 9, 159–176, doi:10.1177/2050640620972602.

92. Gerayeli, F. V.; Milne, S.; Cheung, C.; Li, X.; Yang, C.W.T.; Tam, A.; Choi, L.H.; Bae, A.; Sin, D.D. COPD and the risk of poor outcomes in COVID-19: A systematic review and meta-analysis. EClinicalMedicine 2021, 33, doi:10.1016/j.eclinm.2021.100789.

93. Almansa, R.; Socias, L.; Sanchez-Garcia, M.; Martín-Loeches, I.; Del Olmo, M.; Andaluz-Ojeda, D.; Bobillo, F.; Rico, L.; Herrero, A.; Roig, V.; et al. Critical COPD respiratory illness is linked to increased transcriptomic activity of neutrophil proteases genes. BMC Res. Notes 2012, 5, 1–8, doi:10.1186/1756-0500-5-401.

94. Sng, J.H.J.; Prazakova, S.; Thomas, P.S.; Herbert, C. MMP-8, MMP-9 and Neutrophil Elastase in Peripheral Blood and Exhaled Breath Condensate in COPD. COPD J. Chronic Obstr. Pulm. Dis. 2017, 14, 238–244, doi:10.1080/15412555.2016.1249790.

95. Gao, S.; Zhu, H.; Zuo, X.; Luo, H. Cathepsin g and its role in inflammation and autoimmune diseases. Arch. Rheumatol. 2018, 33, 748–749.

96. Asad, S.; Wegler, C.; Ahl, D.; Bergström, C.A.S.; Phillipson, M.; Artursson, P.; Teleki, A. Proteomics-Informed Identification of Luminal Targets For In Situ Diagnosis of Inflammatory Bowel Disease. J. Pharm. Sci. 2021, 110, 239–250, doi:10.1016/j.xphs.2020.11.001.

97. Akgun, E.; Tuzuner, M.B.; Sahin, B.; Kilercik, M.; Kulah, C.; Cakiroglu, H.N.; Serteser, M.; Unsal, I.; Baykal, A.T. Proteins associated with neutrophil degranulation are upregulated in nasopharyngeal swabs from SARS-CoV-2 patients. PLoS One 2020, 15, e0240012, doi:10.1371/journal.pone.0240012.

98. Reusch, N.; De Domenico, E.; Bonaguro, L.; Schulte-Schrepping, J.; Baßler, K.; Schultze, J.L.; Aschenbrenner, A.C. Neutrophils in COVID-19. Front. Immunol. 2021, 12, 952.

99. Metzemaekers, M.; Cambier, S.; Blanter, M.; Vandooren, J.; Carvalho, A.C.; Malengier-Devlies, B.; Vanderbeke, L.; Jacobs, C.; Coenen, S.; Martens, E.; et al. Kinetics of peripheral blood neutrophils in severe coronavirus disease 2019. Clin. Transl. Immunol. 2021, 10, e1271, doi:10.1002/cti2.1271.

100. Meizlish, M.L.; Pine, A.B.; Bishai, J.D.; Goshua, G.; Nadelmann, E.R.; Simonov, M.; Chang, C.H.; Zhang, H.; Shallow, M.; Bahel, P.; et al. A neutrophil activation signature predicts critical illness and mortality in COVID-19. Blood Adv. 2021, 5, 1164–1177, doi:10.1182/bloodadvances.2020003568.

101. Ackermann, M.; Anders, H.-J.; Bilyy, R.; Bowlin, G.L.; Daniel, C.; De Lorenzo, R.; Egeblad, M.; Henneck, T.; Hidalgo, A.; Hoffmann, M.; et al. Patients with COVID-19: in the dark-NETs of neutrophils. Cell Death Differ. 2021, 11, 12, doi:10.1038/s41418-021-00805-z.

102. Albeituni, S.; Verbist, K.C.; Tedrick, P.E.; Tillman, H.; Picarsic, J.; Bassett, R.; Nichols, K.E. Mechanisms of action of ruxolitinib in murine models of hemophagocytic lymphohistiocytosis. Blood 2019, 134, 147–159, doi:10.1182/blood.2019000761.

103. Ding, J.; Hostallero, D.E.; El Khili, M.R.; Fonseca, G.J.; Milette, S.; Noorah, N.; Guay-Belzile, M.; Spicer, J.; Daneshtalab, N.; Sirois, M.; et al. A network-informed analysis of SARS-CoV-2 and hemophagocytic lymphohistiocytosis genes’ interactions points to Neutrophil extracellular traps as mediators of thrombosis in COVID-19. PLoS Comput. Biol. 2021, 17, e1008810, doi:10.1371/journal.pcbi.1008810.

104. Takeshita, S.; Kawamura, Y.; Kanai, T.; Yoshida, Y.; Tsujita, Y.; Nonoyama, S. The Role of Neutrophil Activation in the Pathogenesis of Kawasaki Disease. Pediatr. Infect. Dis. Open Access 2017, 3, 1, doi:10.21767/2573-0282.100057.

105. Biezeveld, M.H.; Van Mierlo, G.; Lutter, R.; Kuipers, I.M.; Dekker, T.; Hack, C.E.; Newburger, J.W.; Kuijpers, T.W. Sustained activation of neutrophils in the course of Kawasaki disease: an association with matrix metalloproteinases. Clin. Exp. Immunol. 2005, 141, 183–188, doi:10.1111/J.1365-2249.2005.02829.X.

106. Hu, J.; Qian, W.; Yu, Z.; Xu, T.; Ju, L.; Hua, Q.; Wang, Y.; Ling, J.J.; Lv, H. Increased Neutrophil Respiratory Burst Predicts the Risk of Coronary Artery Lesion in Kawasaki Disease. Front. Pediatr. 2020, 8, doi:10.3389/FPED.2020.00391.

107. Balamayooran, G.; Batra, S.; Fessler, M.B.; Happel, K.I.; Jeyaseelan, S. Mechanisms of Neutrophil Accumulation in the Lungs Against Bacteria. Am. J. Respir. Cell Mol. Biol. 2010, 43, 5, doi:10.1165/RCMB.2009-0047TR.

108. Margraf, A.; Ley, K.; Zarbock, A. Neutrophil Recruitment: From Model Systems to Tissue-Specific Patterns. Trends Immunol. 2019, 40, 613–634.

109. Mortaz, E.; Alipoor, S.D.; Adcock, I.M.; Mumby, S.; Koenderman, L. Update on neutrophil function in severe inflammation. Front. Immunol. 2018, 9, 2171.

110. Delorey, T.M.; Ziegler, C.G.K.; Heimberg, G.; Normand, R.; Yang, Y.; Segerstolpe, Å.; Abbondanza, D.; Fleming, S.J.; Subramanian, A.; Montoro, D.T.; et al. COVID-19 tissue atlases reveal SARS-CoV-2 pathology and cellular targets. Nature 2021, doi:10.1038/s41586-021-03570-8.

111. Andonegui-Elguera, S.; Taniguchi-Ponciano, K.; Gonzalez-Bonilla, C.R.; Torres, J.; Mayani, H.; Herrera, L.A.; Peña-Martínez, E.; Silva-Román, G.; Vela-Patiño, S.; Ferreira-Hermosillo, A.; et al. Molecular Alterations Prompted by SARS-CoV-2 Infection: Induction of Hyaluronan, Glycosaminoglycan and Mucopolysaccharide Metabolism. Arch. Med. Res. 2020, 51, 645–653, doi:10.1016/j.arcmed.2020.06.011.

112. Li, Y.; Duche, A.; Sayer, M.R.; Roosan, D.; Khalafalla, F.G.; Ostrom, R.S.; Totonchy, J.; Roosan, M.R. SARS-CoV-2 early infection signature identified potential key infection mechanisms and drug targets. BMC Genomics 2021, 22, doi:10.1186/s12864-021-07433-4.

113. Gardinassi, L.G.; Souza, C.O.S.; Sales-Campos, H.; Fonseca, S.G. Immune and Metabolic Signatures of COVID-19 Revealed by Transcriptomics Data Reuse. Front. Immunol. 2020, 11, doi:10.3389/fimmu.2020.01636.

114. Lucas, C.; Wong, P.; Klein, J.; Castro, T.B.R.; Silva, J.; Sundaram, M.; Ellingson, M.K.; Mao, T.; Oh, J.E.; Israelow, B.; et al. Longitudinal analyses reveal immunological misfiring in severe COVID-19. Nature 2020, 584, 463–469, doi:10.1038/s41586-020-2588-y.

115. Protasio Veras, F.; Pontelli, M.; Silva, C.; Toller-Kawahisa, J.; de Lima, M.; Nascimento, D.; Schneider, A.; Caetité, D.; Tavares, L.; Paiva, I.; et al. SARS-CoV-2-triggered neutrophil extracellular traps mediate COVID-19 pathology. J. Exp. Med. 2020, 217, doi:10.1084/JEM.20201129.

116. Derakhshani, A.; Hemmat, N.; Asadzadeh, Z.; Ghaseminia, M.; Shadbad, M.A.; Jadideslam, G.; Silvestris, N.; Racanelli, V.; Baradaran, B. Arginase 1 (Arg1) as an Up-Regulated Gene in COVID-19 Patients: A Promising Marker in COVID-19 Immunopathy. J. Clin. Med. 2021, 10, 1051, doi:10.3390/JCM10051051.

117. Agresti, N.; Lalezari, J.P.; Amodeo, P.P.; Mody, K.; Mosher, S.F.; Seethamraju, H.; Kelly, S.A.; Pourhassan, N.Z.; Sudduth, C.D.; Bovinet, C.; et al. Disruption of CCR5 signaling to treat COVID-19-associated cytokine storm: Case series of four critically ill patients treated with leronlimab. J. Transl. Autoimmun. 2021, 4, 100083.

118. Guaraldi, G.; Meschiari, M.; Cozzi-Lepri, A.; Milic, J.; Tonelli, R.; Menozzi, M.; Franceschini, E.; Cuomo, G.; Orlando, G.; Borghi, V.; et al. Tocilizumab in patients with severe COVID-19: a retrospective cohort study. Lancet Rheumatol. 2020, 2, e474–e484, doi:10.1016/S2665-9913(20)30173-9.

119. Robinson, P.C.; Liew, D.F.L.; Liew, J.W.; Monaco, C.; Richards, D.; Shivakumar, S.; Tanner, H.L.; Feldmann, M. The Potential for Repurposing Anti-TNF as a Therapy for the Treatment of COVID-19. Med 2020, 1, 90–102, doi:10.1016/j.medj.2020.11.005.

120. Huet, T.; Beaussier, H.; Voisin, O.; Jouveshomme, S.; Dauriat, G.; Lazareth, I.; Sacco, E.; Naccache, J.M.; Bézie, Y.; Laplanche, S.; et al. Anakinra for severe forms of COVID-19: a cohort study. Lancet Rheumatol. 2020, 2, e393–e400, doi:10.1016/S2665-9913(20)30164-8.

121. Gozzetti, A.; Capochiani, E.; Bocchia, M. The Janus kinase 1/2 inhibitor ruxolitinib in COVID-19. Leukemia 2020, 34, 2815–2816.

122. Rizk, J.G.; Kalantar-Zadeh, K.; Mehra, M.R.; Lavie, C.J.; Rizk, Y.; Forthal, D.N. Pharmaco-Immunomodulatory Therapy in COVID-19. Drugs 2020, 80, 1267–1292, doi:10.1007/s40265-020-01367-z.

123. Chiang, C.C.; Korinek, M.; Cheng, W.J.; Hwang, T.L. Targeting Neutrophils to Treat Acute Respiratory Distress Syndrome in Coronavirus Disease. Front. Pharmacol. 2020, 11, 1576.

124. Korkmaz, B.; Lesner, A.; Marchand-Adam, S.; Moss, C.; Jenne, D.E. Lung Protection by Cathepsin C Inhibition: A New Hope for COVID-19 and ARDS? J. Med. Chem. 2020, 63, 13258–13265.

125. Dennison, D.; Al Khabori, M.; Al Mamari, S.; Aurelio, A.; Al Hinai, H.; Al Maamari, K.; Alshekaili, J.; Al Khadouri, G. Circulating activated neutrophils in COVID-19: An independent predictor for mechanical ventilation and death. Int. J. Infect. Dis. 2021, 106, 155–159, doi:10.1016/J.IJID.2021.03.066.

126. Seery, V.; Raiden, S.C.; Algieri, S.C.; Grisolía, N.A.; Filippo, D.; De Carli, N.; Di Lalla, S.; Cairoli, H.; Chiolo, M.J.; Meregalli, C.N.; et al. Blood neutrophils from children with COVID-19 exhibit both inflammatory and anti-inflammatory markers. EBioMedicine 2021, 67, 103357, doi:10.1016/J.EBIOM.2021.103357/ATTACHMENT/7BCA7452-0548-429A-928A-80540165D002/MMC4.DOCX.

127. Gu, R.; Mao, T.; Lu, Q.; Tianjiao Su, T.; Wang, J. Myeloid dysregulation and therapeutic intervention in COVID-19. Semin. Immunol. 2021, 55, 101524, doi:10.1016/J.SMIM.2021.101524.

