## Supplementary material for "Severe COVID-19 shares a common neutrophil activation signature with other acute inflammatory states": Key resource table Methods LF Schimke 2022.docx

**KEY RESOURCES TABLE**

| **REAGENT or RESOURCE** | **SOURCE** | **IDENTIFIER** |
| --- | --- | --- |
| **Deposited data** | | |
| mircoarray data of HLH patients and healthy controls | Sumegi et al. 2011 | GEO: GSE26050 |
| bulk-RNA seq data of COVID-19 patients and controls | Arunachalam et al. 2020 | GEO: GSE152418 |
| bulk-RNA seq data of COVID-19 patients and controls | Liebermann et al. 2020 | GEO: GSE152075 |
| bulk-RNA seq data of COVID-19 patients and patients with other respiratory diseases | Mick et al. 2020 | GEO: GSE156063 |
| bulk-RNA seq data of COVID-19 patients and controls | Ng et al. 2021 | GEO: GSE163151 |
| bulk-RNA seq data of COVID-19 patients and controls | Thair et al. 2021 | GEO: GSE152641 |
| bulk-RNA seq data of COVID-19 patients and patients with other respiratory symptoms | Overmyer et al. 2020 | GEO: GSE157103 |
| scRNA seq data of COVID-19 patients and controls | Schulte-Schrepping et al. 2020 | EGA: EGAS00001004571 |
| bulk-RNA seq data of COVID-19 patients, healthy controls, and patients with Influenza, bacterial pneumonia, and seasonal Coronavirus other than SARS-CoV-2 | McClain et al.2021 | GSE161731 |
| bulk-RNA seq data of patients with multisystem inflammatory syndrome in children (MIS-C) and healthy controls | Beckmann et al. 2021 | GSE178388 |
| microarray data of patients with Kawasaky disease (KD) and healthy controls | Wright et al. 2018 | GSE73461 |
| **Software and algorithms** | | |
| NetworkAnalyst 3.0 | Zhou, G. et al. 2019 | https://www.networkanalyst.ca/ |
| Limma voom pipeline | Law, C. W. et al. 2014 | https://doi.org/10.1186/gb-2014-15-2-r29 |
| jvenn | Bardou, P. et al. 2014 | http://jvenn.toulouse.inra.fr/app/example.html |
| Circos online tool | Krzywinski, M. et al. 2009 | http://mkweb.bcgsc.ca/tableviewer/ |
| Seurat V.4 | Hao, Y. et al. 2020 | https://doi.org/10.1016/j.cell.2021.04.048https://satijalab.org/seurat/ |
| NAViGaTOR 3.0.14 | Brown, K. R. et al. 2009 | doi:10.1093/bioinformatics/btp595 |
| Integrated Interactions Database, IID version 2020-05 | Kotlyar M. et al. 2018 | http://ophid.utoronto.ca/iid |
| ClusterProfiler | Yu, G et al. 2012 | doi:10.1089/omi.2011.0118. |
| Enrichr | Chen, E. Y. et al. 2013; Kuleshov, M. V. et al.2016 | https://maayanlab.cloud/Enrichr/ |
| Morpheus | Starruß, J. et al. 2014 | https://software.broadinstitute.org/morpheus/ |
| R version 4.0.5 | R Core Team (2020) | https://www.r-project.org/ |
| R studio Version 1.4.1106 | RStudio Team (2020) | http://www.rstudio.com/. |
| circlize R package | Zuguang Gu et al. 2020 | https://github.com/jokergoo/circlize |
| ggpubr R package | Alboukadel Kassambara, 2020 | https://rpkgs.datanovia.com/ggpubr/ |
| lemon R package | Stefan McKinnon Edwards et al. 2020 | https://github.com/stefanedwards/lemon |
| ggplot2 R package | Wickham H, 2016 | https://ggplot2.tidyverse.org |
| factoextra R package for PCA analysis | Alboukadel Kassambara and Fabian Mundt, 2017 | https://rpkgs.datanovia.com/factoextra/index.html |
| Canonical Correlation Analysis (CCA) R package | Jendoubi, T. & Strimmer, K., 2019 | https://cran.r-project.org/web/packages/whitening/index.html http://www.strimmerlab.org/software/whitening/ |
| corrgram R package | Kevin Wright, 2021 | https://kwstat.github.io/corrgram/ |
| psych R package | William Revelle, 2021 | https://personality- project.org/r/psych/ |
| inlmisc R package | Jason C Fisher, 2021 | https://github.com/USGS-R/inlmisc |
| tydiverse R package | Hadley Wickham, 2021 | https://github.com/tidyverse/tidyverse/ |
| viridis R package | Simon Garnier et al. 2021 | https://sjmgarnier.github.io/viridis/ |
| randomForest (version 4.6.14) | Andy Liaw and Matthew Wiener 2002 | https://cran.r-project.org/doc/Rnews/Rnews_2002-3.pdf |
| Fisher´s method | Fisher, R. A., 1932 | Fisher, R. A. Statistical Methods for Research Workers 4th edn. (Oliver and Boyd, 1932). |
| nonparametric MANOVA: nparcomp R package | Konietschke, F. et al., 2015 | doi:10.18637/jss.v064.i09 |
| nonparametric Inference for Multivariate Data: npmv R package | Burchett, W. et al., 2017 | doi:10.18637/jss.v076.i04 |
