## Supplementary material for "Severe COVID-19 shares a common neutrophil activation signature with other acute inflammatory states": SupplementalFigures and Legends_07 Feb 2022 LSchimke.pptx

### Slide 1
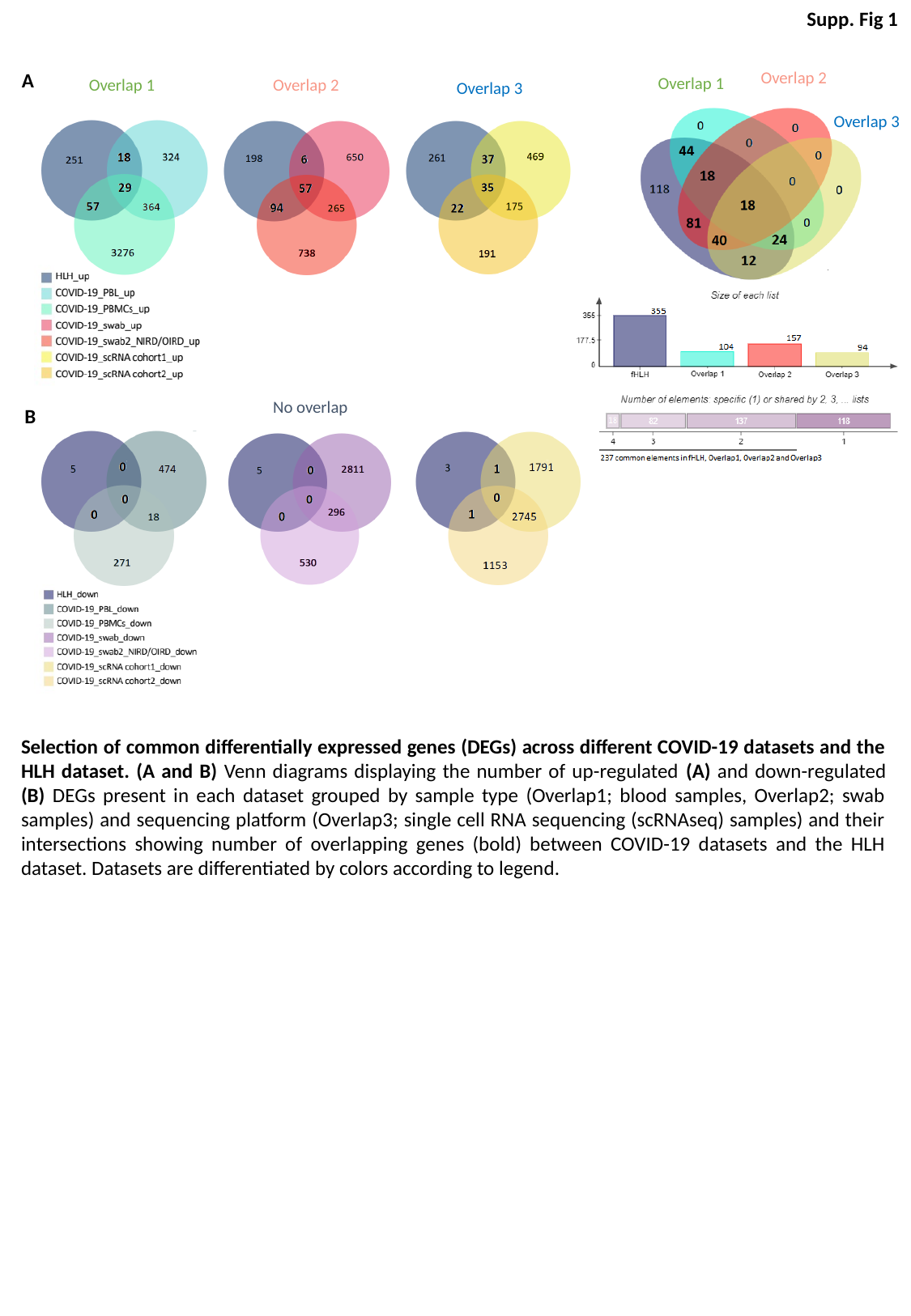

Supp. Fig 1
Overlap 2
A
Overlap 1
Overlap 1
Overlap 2
Overlap 3
Overlap 3
No overlap
B
Selection of common differentially expressed genes (DEGs) across different COVID-19 datasets and the HLH dataset. (A and B) Venn diagrams displaying the number of up-regulated (A) and down-regulated (B) DEGs present in each dataset grouped by sample type (Overlap1; blood samples, Overlap2; swab samples) and sequencing platform (Overlap3; single cell RNA sequencing (scRNAseq) samples) and their intersections showing number of overlapping genes (bold) between COVID-19 datasets and the HLH dataset. Datasets are differentiated by colors according to legend.

### Slide 2
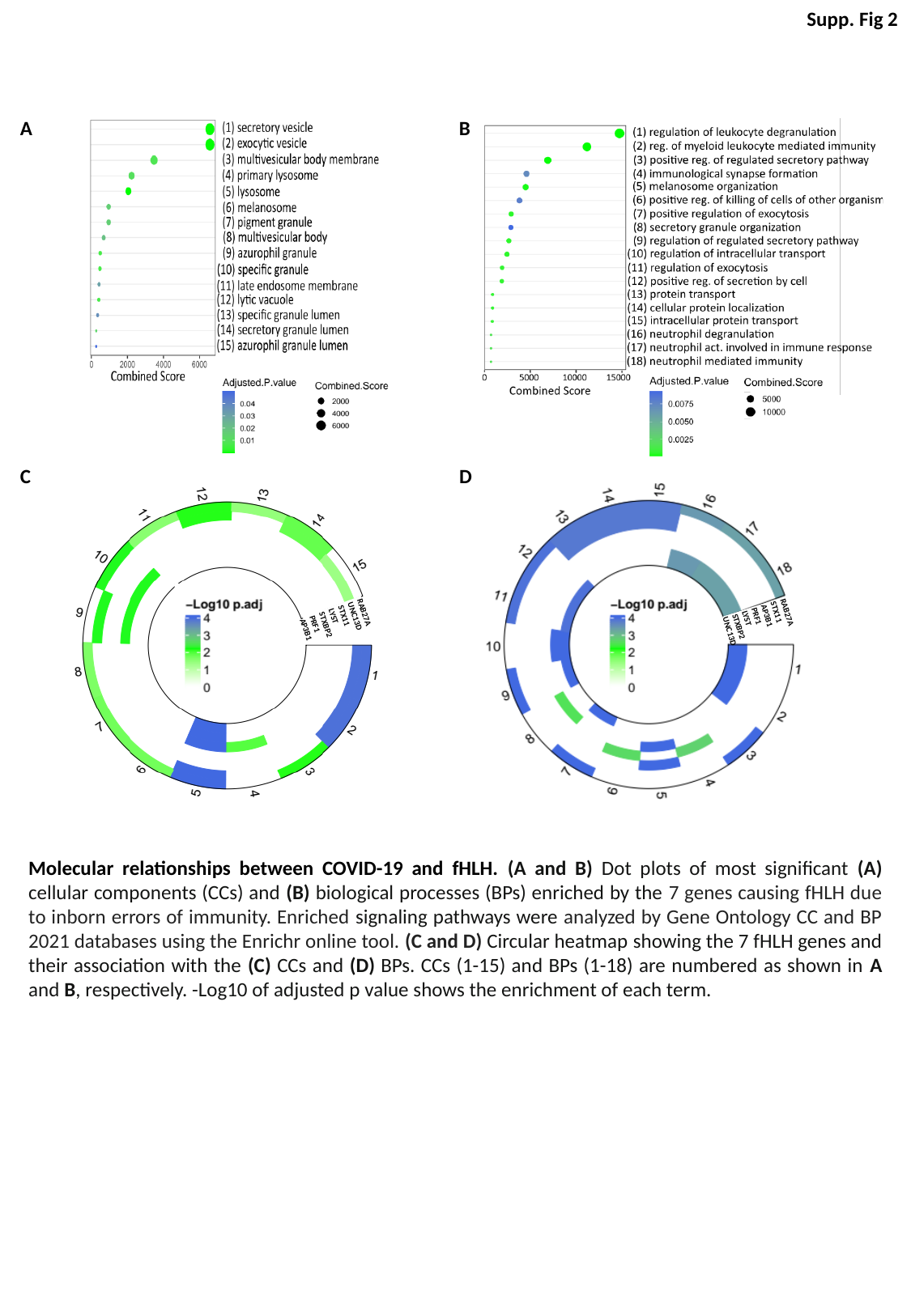

Supp. Fig 2
A
B
C
D
RAB27A
STX11
AP3B1
PRF1
LYST
STXBP2
UNC13D
RAB27A
UNC13D
STX11
LYST
STXBP2
PRF1
AP3B1
Molecular relationships between COVID-19 and fHLH. (A and B) Dot plots of most significant (A) cellular components (CCs) and (B) biological processes (BPs) enriched by the 7 genes causing fHLH due to inborn errors of immunity. Enriched signaling pathways were analyzed by Gene Ontology CC and BP 2021 databases using the Enrichr online tool. (C and D) Circular heatmap showing the 7 fHLH genes and their association with the (C) CCs and (D) BPs. CCs (1-15) and BPs (1-18) are numbered as shown in A and B, respectively. -Log10 of adjusted p value shows the enrichment of each term.

### Slide 3
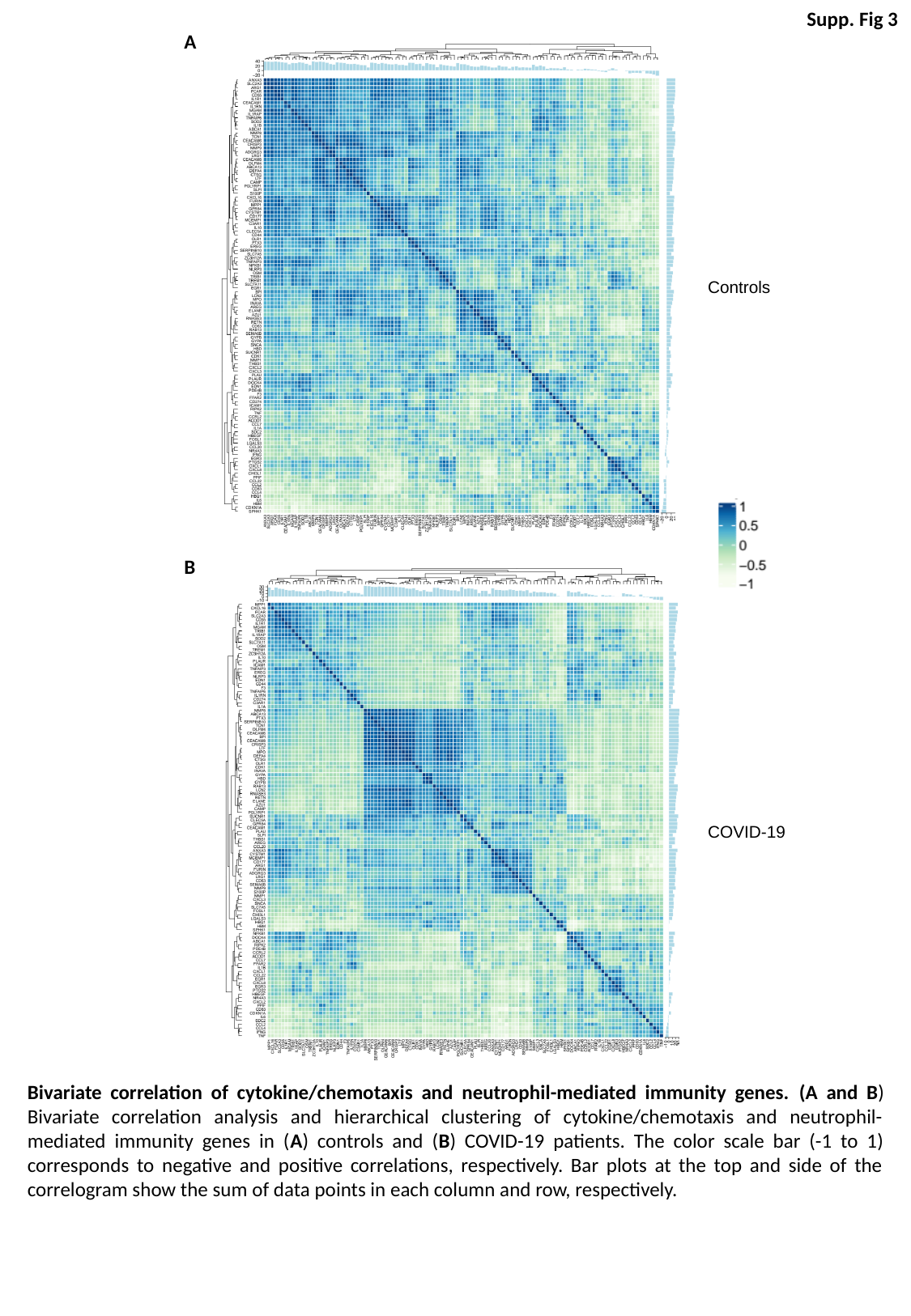

Supp. Fig 3
A
Controls
B
COVID-19
Bivariate correlation of cytokine/chemotaxis and neutrophil-mediated immunity genes. (A and B) Bivariate correlation analysis and hierarchical clustering of cytokine/chemotaxis and neutrophil-mediated immunity genes in (A) controls and (B) COVID-19 patients. The color scale bar (-1 to 1) corresponds to negative and positive correlations, respectively. Bar plots at the top and side of the correlogram show the sum of data points in each column and row, respectively.

### Slide 4
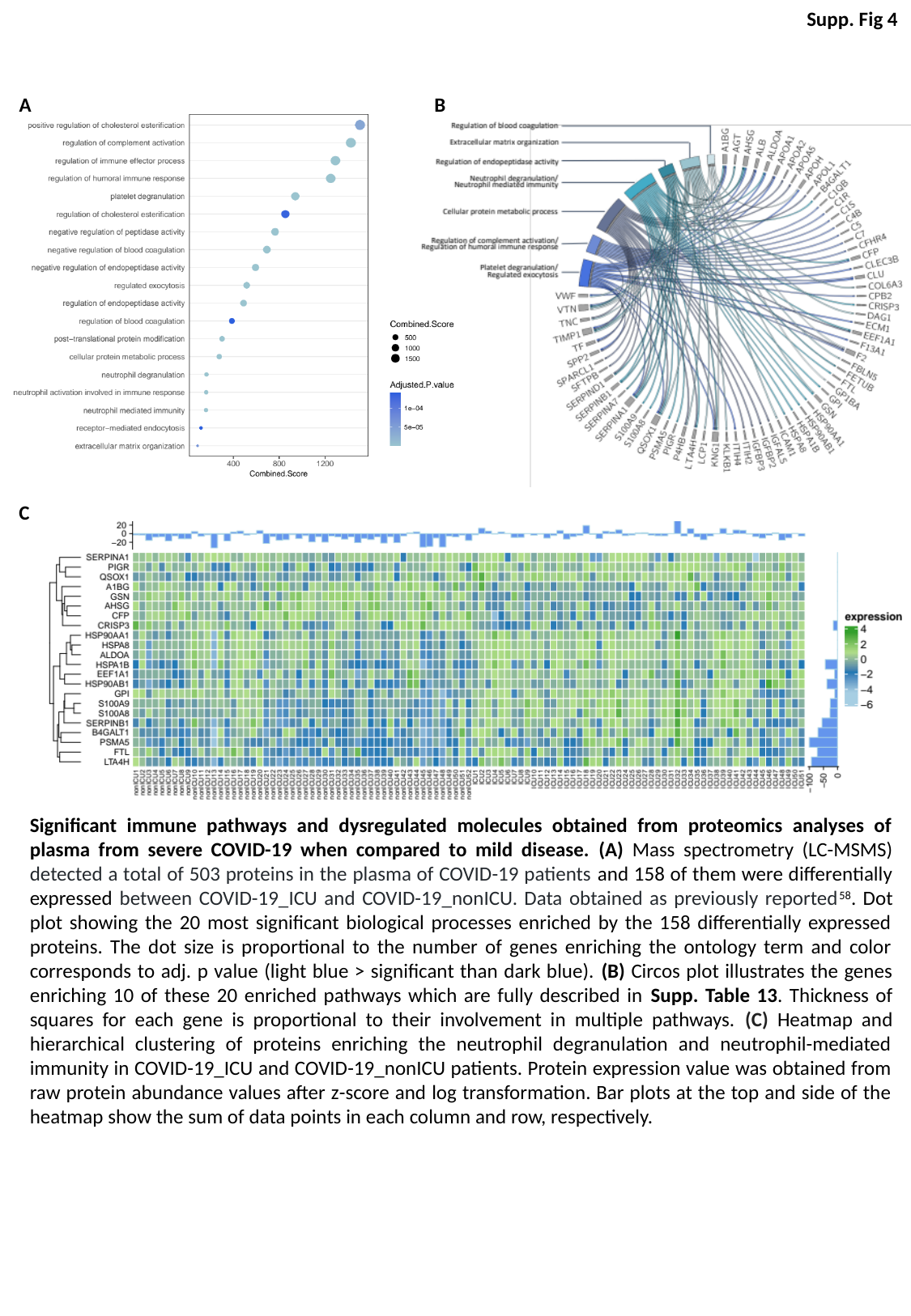

Supp. Fig 4
A
B
C
Significant immune pathways and dysregulated molecules obtained from proteomics analyses of plasma from severe COVID-19 when compared to mild disease. (A) Mass spectrometry (LC-MSMS) detected a total of 503 proteins in the plasma of COVID-19 patients and 158 of them were differentially expressed between COVID-19_ICU and COVID-19_nonICU. Data obtained as previously reported^58^. Dot plot showing the 20 most significant biological processes enriched by the 158 differentially expressed proteins. The dot size is proportional to the number of genes enriching the ontology term and color corresponds to adj. p value (light blue > significant than dark blue). (B) Circos plot illustrates the genes enriching 10 of these 20 enriched pathways which are fully described in Supp. Table 13. Thickness of squares for each gene is proportional to their involvement in multiple pathways. (C) Heatmap and hierarchical clustering of proteins enriching the neutrophil degranulation and neutrophil-mediated immunity in COVID-19_ICU and COVID-19_nonICU patients. Protein expression value was obtained from raw protein abundance values after z-score and log transformation. Bar plots at the top and side of the heatmap show the sum of data points in each column and row, respectively.

### Slide 5
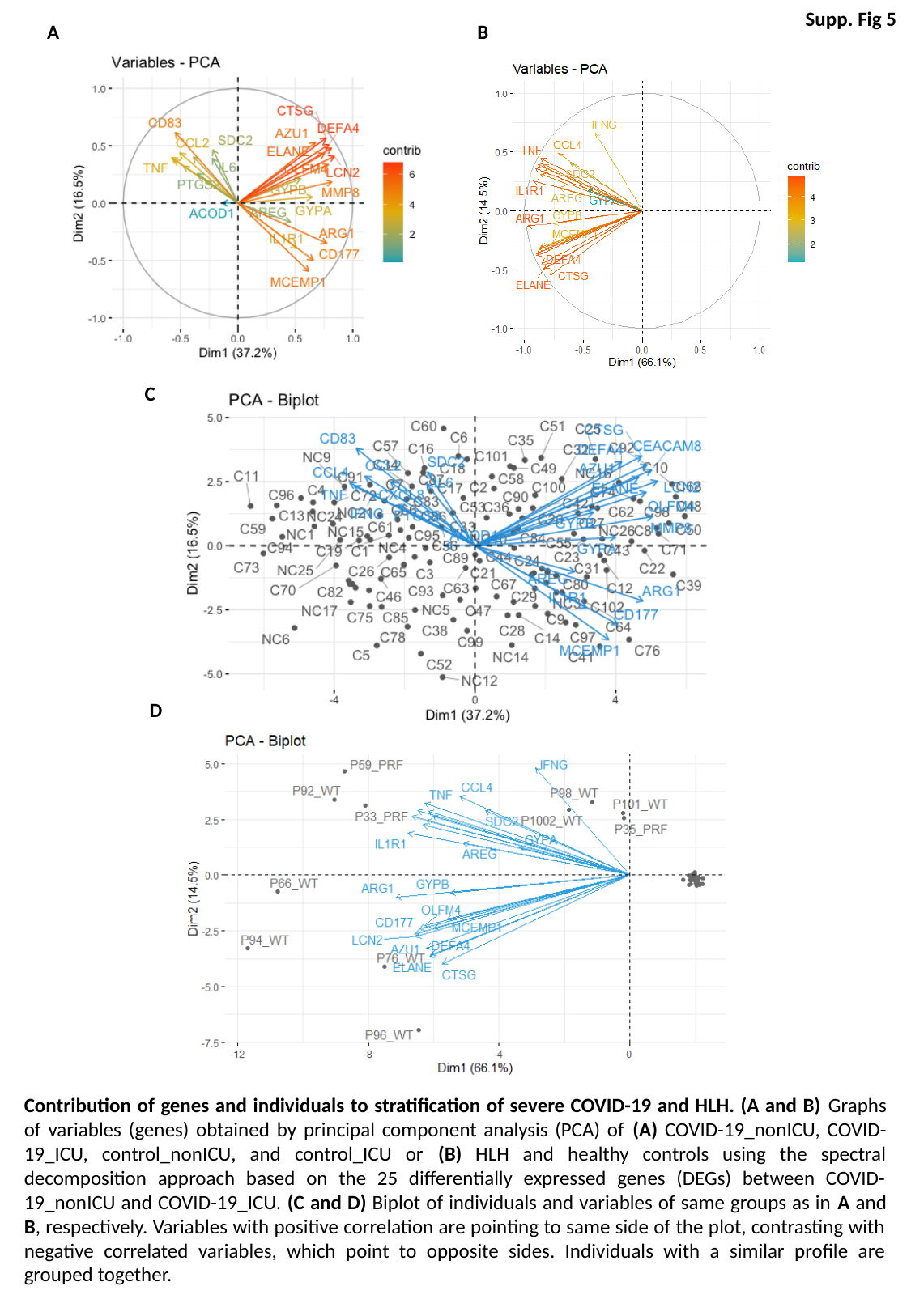

Supp. Fig 5
A
B
C
D
Contribution of genes and individuals to stratification of severe COVID-19 and HLH. (A and B) Graphs of variables (genes) obtained by principal component analysis (PCA) of (A) COVID-19_nonICU, COVID-19_ICU, control_nonICU, and control_ICU or (B) HLH and healthy controls using the spectral decomposition approach based on the 25 differentially expressed genes (DEGs) between COVID-19_nonICU and COVID-19_ICU. (C and D) Biplot of individuals and variables of same groups as in A and B, respectively. Variables with positive correlation are pointing to same side of the plot, contrasting with negative correlated variables, which point to opposite sides. Individuals with a similar profile are grouped together.

### Slide 6
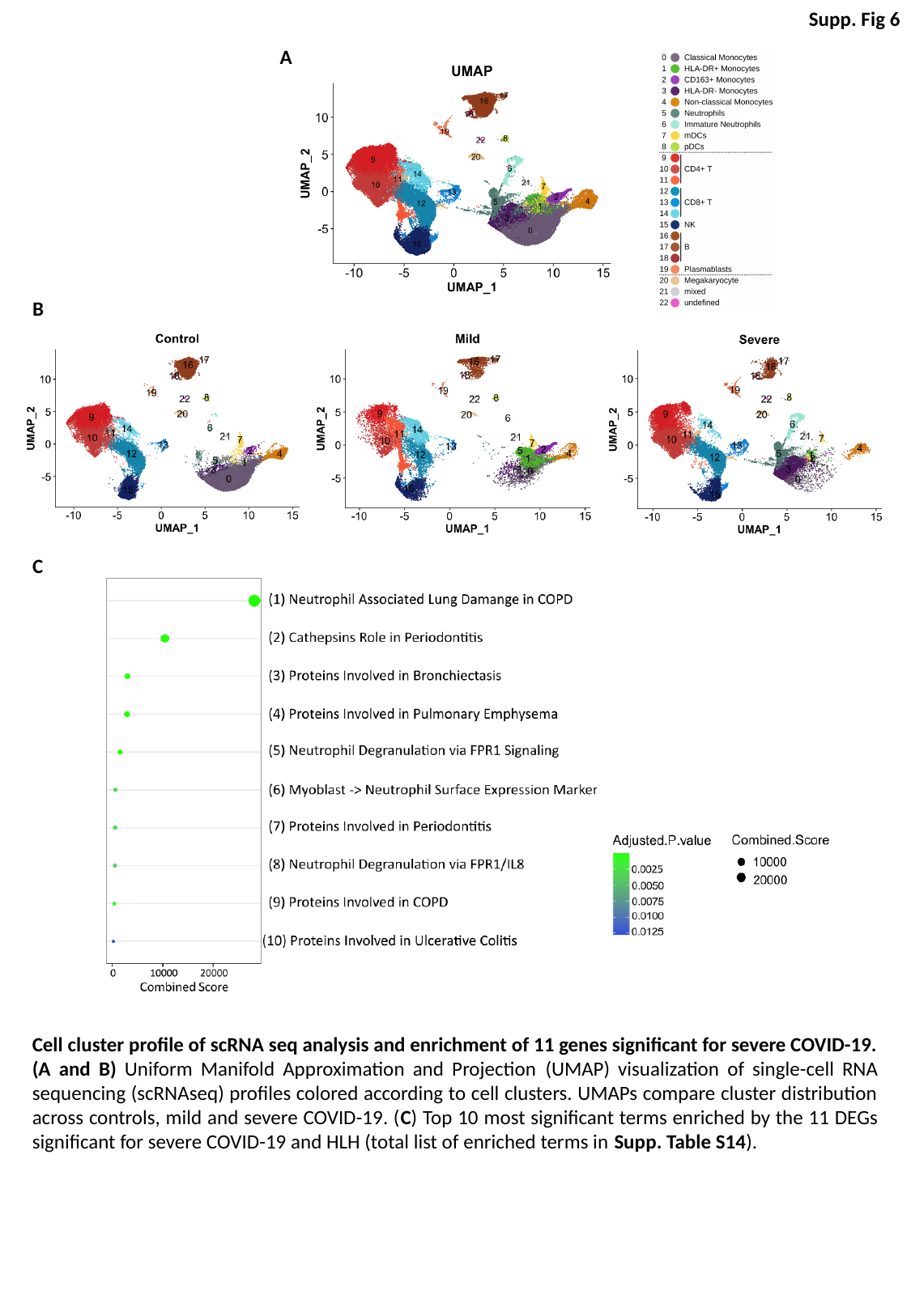

Supp. Fig 6
A
B
C
Cell cluster profile of scRNA seq analysis and enrichment of 11 genes significant for severe COVID-19. (A and B) Uniform Manifold Approximation and Projection (UMAP) visualization of single-cell RNA sequencing (scRNAseq) profiles colored according to cell clusters. UMAPs compare cluster distribution across controls, mild and severe COVID-19. (C) Top 10 most significant terms enriched by the 11 DEGs significant for severe COVID-19 and HLH (total list of enriched terms in Supp. Table S14).
